# Identification of a C3b-specific nanobody that does not bind C3 and blocks alternative pathway convertases

**DOI:** 10.64898/2026.08.30.748118

**Authors:** Eva M Struijf, Bart W. Bardoel, Joanne E. van Keulen, Maartje Ruyken, Danique Y. Siere, Douwe J. Dijkstra, Gillian Dekkers, Piet Gros, Raimond Heukers, Dani A.C. Heesterbeek, Suzan H.M. Rooijakkers, Jamie S. Depelteau

**Affiliations:** Medical Microbiology, University Medical Center Utrecht, Utrecht University, Utrecht, the Netherlands; Structural Biochemistry, Bijvoet Centre for Biomolecular Research, Department of Chemistry, Faculty of Science, Utrecht University, Utrecht, the Netherlands; QVQ Holding BV, Utrecht, the Netherlands

**Keywords:** Innate immunity, complement system, nanobody, C3, C3b, alternative pathway, convertase, inhibition

## Abstract

The human complement system is a protein network in blood and other body fluids that fights invading pathogens and is involved in maintaining homeostasis. A central step in the complement cascade is the conversion of complement protein C3 to C3b by convertase enzymes. Upon cleavage of C3, the nascent C3b molecule undergoes a large conformational change which exposes new epitopes. Molecules that discriminate between C3b and C3 could provide a powerful tool to selectively bind surface-bound complement activation products while leaving the circulating precursors untouched. In this study, we developed C3b-specific nanobodies by generating phage libraries from llamas immunized with purified and surface-bound C3b. We describe UNbC3b-1 as a high-affinity binder that recognizes soluble and surface-bound C3b molecules but does not bind to C3 in its native state. A 4.2 Å structure of the UNbC3b-1:C3b complex generated by cryogenic electron microscopy shows that UNbC3b-1 binds C3b at the interface of MG3, MG4, and MG6, overlapping with the complement receptor immunoglobulin (CRIg) binding site. Functionally, we show that UNbC3b-1 inhibits complement activity in the alternative pathway (AP), likely by preventing association between substrate C3 and the AP C3 convertase (C3bBb). Altogether, nanobody UNbC3b-1 is a valuable tool to specifically bind C3b molecules and to selectively inhibit the AP convertases of the complement system.

## INTRODUCTION

The complement system is a large protein network in blood that is part of the human innate immune system. It plays a crucial role in the fight against invading microbes and it is involved in maintaining homeostasis by the removal of apoptotic cells and immune complexes (1, 2). Activation of the classical pathway (CP) and lectin pathway (LP) initiates a proteolytic cascade (involving complement proteins C1, C2 and C4) on the target cell surface, which results in the formation of the CP/LP C3 convertase, C4b2b (1, 2). The alternative pathway (AP) is initiated when C3b molecules are formed via the CP and LP, or when C3 spontaneously hydrolyzes into the C3b-like molecule, C3(H_2_O). Together with C3b, or C3(H_2_O), complement components Factor B (FB) and Factor D (FD) can form the AP C3 convertase, C3bBb (1, 2). Cleavage of C3 by convertases initiates the formation of pro-inflammatory chemoattractants (C3a and C5a), opsonins (e.g. C3b and iC3b), and membrane attack complexes (MAC) that can directly lyse target cells.

The cleavage of C3 into C3b exposes hidden features that are not present in native C3. C3 is a large protein (185 kDa) comprised of an alpha chain (110 kDa) and a beta chain (75 kDa) that are covalently bound to each other (3, 4). When the convertase cleaves C3, the ANA domain (C3a, 9 kDa) located near the center of the molecule is released (5, 6). The remaining C3b molecule (176 kDa) undergoes major conformational rearrangements in which the CUB and TED domains make the largest movements, exposing the reactive thioester with which C3b attaches to a surface (**Fig. 1A**). Deposited C3b functions as an opsonin by helping phagocytic cells to recognize and engulf target cells via complement receptors 1 (CR1), CR3, and CRIg (CR of immunoglobulin) (1, 2). Furthermore, newly deposited C3b molecules induce an amplification loop when they interact with FB and FD, forming new AP C3 convertases that convert other C3 molecules. When the deposition of C3b molecules reaches a high density, C3 convertases switch substrate from C3 to C5, forming C5 convertases (CP/LP: C4b2bC3b, AP: C3bBbC3b) (7–10). Newly formed C5 convertases cleave circulating complement component C5 into the anaphylatoxin C5a, and the activation product C5b, which initiates the formation of the target cell lysing MAC (C5b-9).

**Figure 1:**
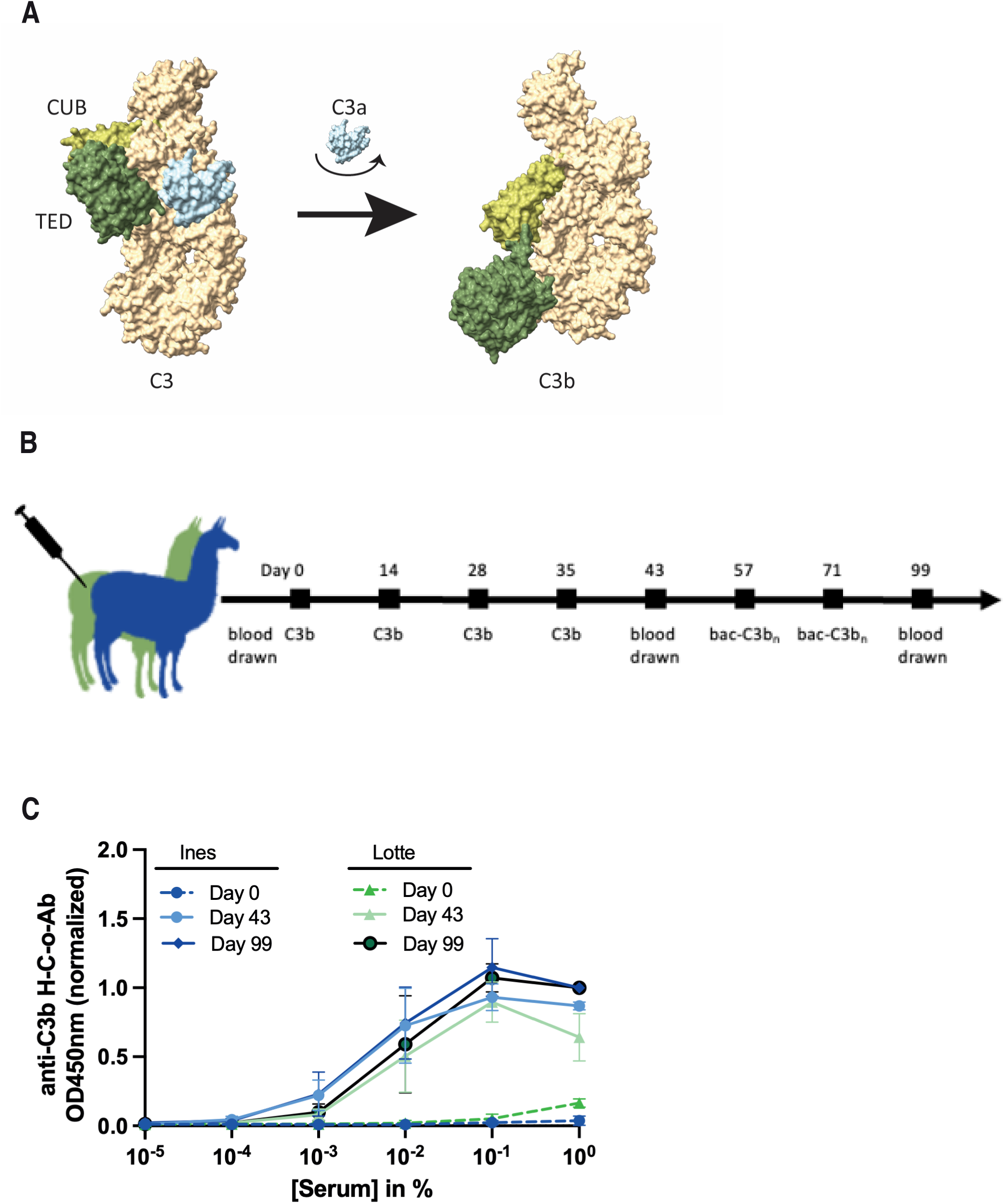
Successful immunization of two llamas with purified human C3b. (A) Schematic presentation of structural rearrangements upon C3 cleavage into C3a (ANA, light blue) and C3b (beige and green), using the published structures of C3 (PDB 2A73) and C3b (PDB 2I07) and depicted in a space filling representation. The CUB (light green) and TED (dark green) domains are highlighted. (B) Immunization scheme of the two llamas, immunized with purified human C3b and bacteria opsonized with high densities of C3b molecules (bac-C3b_n_). (C) Detection of anti-C3b antibodies in llama serum before immunizations (day 0), after immunizations with purified C3b in solution (day 43) and after immunizations with C3b-opsonized bacteria (day 99). C3b-binding llama antibodies were detected in an ELISA setup using polyclonal rabbit-anti-nanobody QE19 antibodies and donkey-anti-rabbit-HRP antibodies, at an OD of 450 nm. Data represents mean ± SD of two individual experiments.

Although the complement system is tightly regulated to prevent uncontrolled activation, unwanted overactivation is associated with a wide variety of inflammatory and autoimmune diseases, including paroxysmal nocturnal hemoglobinuria (PNH), C3 glomerulopathy (C3G), age-related macular degeneration (AMD), rheumatoid arthritis, and many other diseases (11–13). Therefore, significant research efforts have been made to develop clinically relevant drugs that target specific pathways and proteins within the system. In the last decades, a wide variety of C3-targeting molecules, monoclonal antibodies and nanobodies have been developed to inhibit complement activity (14). The compstatin family, consisting of multiple cyclic peptides that inhibit C3 conversion by binding C3 and C3b, is investigated best (15). Pegcetacoplan (also known as Empavelia®), is a second-generation analog of compstatin, and was recently approved by the FDA as the first (and still only) C3-targeting molecule in the clinic (15, 16). A common feature of most complement inhibitors is that they target circulating non-activated complement precursors. However, more efforts are now directed at developing inhibitors that specifically bind to activated proteins while leaving circulating precursors untouched.

In this study, we describe the development and characterization of a novel llama derived anti-C3b nanobody that specifically recognizes C3b but not precursor C3. Our high affinity nanobody binds at the interface of MG3, MG4, and MG6, which overlaps with the binding site of CRIg. Mechanistic studies show that the nanobody potently inhibits the alternative, but not the classical or lectin pathway by preventing C3 cleavage by AP convertases.

## RESULTS

### Immunization of two llamas with human C3b

To develop C3b-specific nanobodies, two llamas were first immunized with four injections of purified human C3b (**Fig. 1B**). Subsequently, llamas were boosted with two injections containing C3b-opsonized bacteria to steer the llama’s immune response towards C3b-epitopes that are only available when C3b is in the surface-bound orientation. To monitor the immune response and create nanobody phage libraries, blood was drawn at day 0, 43, and 99 to collect serum and peripheral blood mononuclear cells (PBMCs). By incubating C3b-coated microtiter plates with llama sera and detecting antibody binding to C3b, we confirmed that those immunizations generated C3b-binding antibodies at day 43 and 99, that were not present at day 0 (**Fig. 1C**).

### C3b-specific nanobodies identification via phage display

PBMCs collected at day 99 were used to generate two phagemid libraries, each with a size of 10^9^ clones. Selection of C3b-specific phages was carried out using beads coated with high densities of C3b as bait during phage display (16). C3b was biotinylated via its thioester moiety and incubated with magnetic streptavidin beads allowing C3b to bind the bead surface in its natural orientation. To obtain a diverse panel of nanobodies recognizing C3b but not precursor C3, we performed different phage panning strategies (**Table 1**). For example, for strategy #9, phages were incubated with high-density C3b-beads, in the presence of high concentrations C3 and C3b in solution. Next, unbound phages and phages bound to C3 and C3b in solution were removed by extensive washing of the beads (negative selection). Phages bound to high density C3b-beads were subsequently eluted using a high pH buffer (positive selection). Bacterial output libraries were cultured and plated to obtain single colonies.

**Table I:** Selection strategies for phage display.

| # | Library |  | Round 1 |  |  | Round 2 |  |  |
| --- | --- | --- | --- | --- | --- | --- | --- | --- |
|  | Day | Llama | surface | [C3b]* | Negative selection | surface | [C3b] | Negative selection |
| 1 | 43 | 1 & 2 | Plate | 5.0 | none | Plate | 0.1 | C3<br>C5 |
| 2 |  |  |  |  |  |  | 1.0 | none |
| 3 |  |  |  |  |  | Beads | 0.2 | C3<br>C5 |
| 4 |  |  |  |  |  |  | 2.0 | C3<br>C5 |
| 5 | 99 | 1 & 2 | Beads | 4.0 | none | - | - | - |
| 6 |  |  |  |  | LD beads**<br>C3<br>C3b | - | - | - |
| 7 |  |  |  |  |  | Beads | 0.1 | LD beads<br>C3<br>C3b |
| 8 |  |  |  |  |  |  | 0.2 | LD beads<br>C3<br>C3b |
| 9 |  |  |  |  |  |  | 0.8 | LD beads<br>C3<br>C3b |

\* Plate: amount of coated C3b in µg/ml. Beads: volume of C3b-coated beads in µl
\*\* Low density (LD) beads: density of C3b molecules is approximately 16x lower than on regular C3b beads.

A total of 92 randomly picked clones and 4 negative controls were cultured in single wells. 76 individual clones were successfully sequenced, revealing 29 clusters with clonally related nanobodies (>80% amino acid sequence similarity in their complementarity determining region H3 (CDR-H3) (**SFig. 1A**). Most of the clones originated from the IGHV3-3*01 and IGHJ4*01 germline sequences, while a small number originated from IGHV3-53*01 and IGHJ6*06 (based on IMGT database (17, 18)). Half of the clusters (14/29) contained >1 clone per cluster. The largest cluster (#22) contained 20 clones, the second largest cluster (#1) contained 9 clones, followed by a cluster (#15) with 5 clones (**SFig. 1A & B**). These numbers indicate that our phage display strategy enriched for specific CDR-H3 clusters, of which individual clones were assessed for binding to C3b.

### Selection of C3b-specific nanobodies

To validate C3b specificity, nanobodies were expressed in the periplasm of *E. coli* BL21 and the crude periplasmic extracts were used to identify C3b-binding nanobodies. Briefly, C3b- and C3-coated microtiter plates were incubated with peri-Nbs and nanobody binding to both proteins was measured. Next, the ratio C3b:C3 binding was calculated and the clones with the highest ratio were considered interesting to follow-up. Interestingly, >70% of our clones had a C3b:C3-binding ratio >1 (**Fig. 2A**) and 25% had a ratio >1.5 (**Fig. 2A**, zoom-in panel). Next, we compared the sequences of the top 25% and selected a panel of eight clones with diverse CDR1, CDR2 and CDR3 regions, and derived from different clusters, for follow up (**Fig. 2B & SFig. 1A & C**).

**Figure 2:**
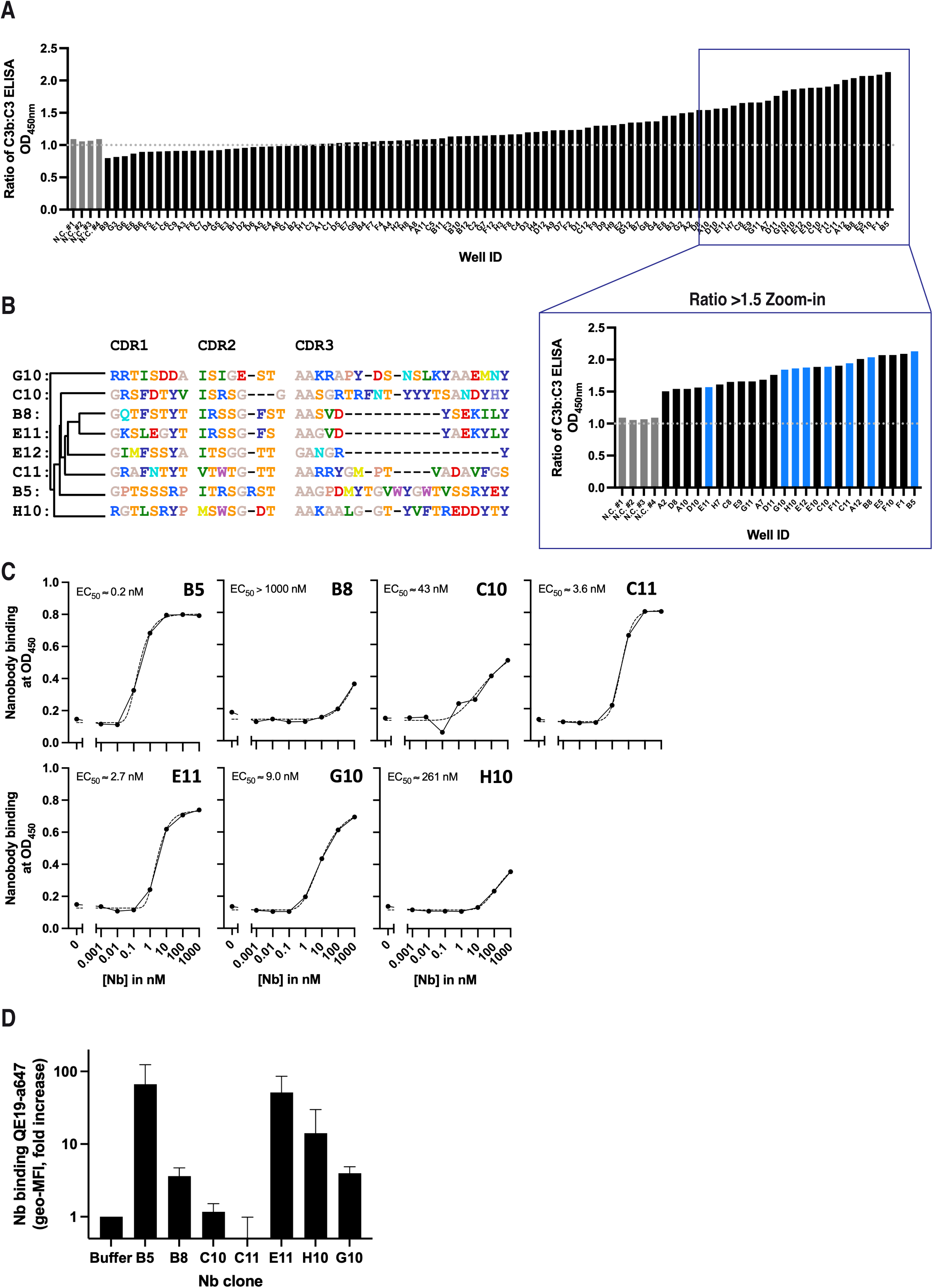
Selection of UNbC3b-1 (clone B5) as specific and inhibitory anti-C3b nanobody. (A) C3b vs C3 binding pattern in ELISA of 92 unpurified periplasmic extracts containing an unknown concentration of nanobody. Negative controls were: #1 periplasmic extract of an irrelevant nanobody; #2 & #4 no bacteria present during culturing; #3 bacteria transformed with empty vector. Microtiter plates coated with C3b and C3 were incubated with periplasmic nanobody extracts (1:5 diluted in PBS), and binding of nanobodies was detected using polyclonal rabbit-anti-VHH QE19 antibodies and donkey-anti-rabbit-HRP antibodies, at an OD of 450 nm. The ratio of C3b:C3 binding was calculated and plotted on the y-axis. Fractions with ratios >1.5 were highlighted in the zoomed-in panel. Blue colored bars indicate clones that were selected to purify and characterize further. Data presented are of one individual experiment. (B) Protein sequence alignment of the eight selected clones from (A). Amino acids annotation is according to the shapely color scheme. (C) Nanobody binding to C3b in ELISA. Microtiter plates coated with randomly oriented C3b were incubated with a concentrations range of purified nanobody (clones B5, B8, C10, C11, E11, G10 and H10). Nanobody binding was detected with polyclonal rabbit-anti-nanobody QE19. To obtain EC50 values (or apparent affinities) binding curves were fitted in GraphPad using the formula “Asymmetric Sigmoidal, 5PL, X is concentration”. Data represents OD450 values of one individual experiment. (D) Nanobody binding to C3b molecules on a bead surface. C3b-beads were incubated with buffer or 100 nM purified nanobody. Nanobody binding was detected using fluorescently labeled (Alexa 647) anti-VHH QE19 and measured by flow cytometry. The fold increase over the background (buffer sample) was calculated and plotted on the y-axis. Data represent mean ± SD of two individual experiments.

Next, seven clones were expressed with a C-terminal Myc- and 6×His-tag in *E. coli* and purified for further characterization. First, we assessed if the purified nanobodies could efficiently bind randomly oriented C3b in ELISA. Clones B5, C11, E11, and G10 bound C3b with an apparent affinity in the low nanomolar range (∼0.2-9.0 nM), while clones B8, C10 and H10 bound C3b with apparent affinities of >40 nM (**Fig. 2C**). Next, we assessed if the nanobodies could also recognize C3b in a surface-bound orientation. To do so, nanobodies were incubated with C3b-beads as described above. Except for C10 and C11, all clones recognized surface-bound C3b to some extent (**Fig. 2D**). Based on the binding to C3b in ELISA (highest C3b:C3 ratio and highest affinity) and on beads, we selected clone B5 to further investigate. From now on, we will refer to this clone (B5) as UNbC3b-1.

### UNbC3b-1 specifically binds C3b with high affinity

Next, we wanted to confirm that UNbC3b-1 is a C3b-specific nanobody that does not bind to C3 and other complement proteins. First, we tested the effect of adding soluble C3b or C3 as competitors for nanobody binding to C3b labelled beads. We observed that UNbC3b-1 binding to C3b-beads is only decreased when C3b is added in high concentrations (**Fig. 3A**). We next determined that UNbC3b-1 also bound to cleavage products of C3b, iC3b and C3c (the larger C3 fragments) but not to the smaller fragments C3dg, C3d, C3a and C3a-desArg in ELISA (**Fig. S2A**). This suggests that UNbC3b-1 does not bind to the TED and ANA domains of C3b. To further assess specificity, we performed an ELISA in which microtiter plates were coated with complement proteins Bb, C4, C4b, C5, C5a, C5b6, and C6 and nanobody binding was measured. UNbC3b-1 did not cross-react with any of the other complement proteins **(Fig. S2B)**. When we coated microtiter plates with C3b and C3, UNbC3b-1 also bound to C3 in several repeats of the experiment, albeit with lower apparent affinities than measured for C3b (**Fig. S2C**). This could indicate that UNbC3b-1, in absence of competitive pressure from C3b, binds to precursor C3. However, since C3 is easily hydrolysed when in contact with a surface (19), the observed binding could be caused by the nanobody binding to the hydrolysed form of C3 (C3(H_2_O)), which is known to adopt a C3b-like conformation (20).

**Figure 3:**
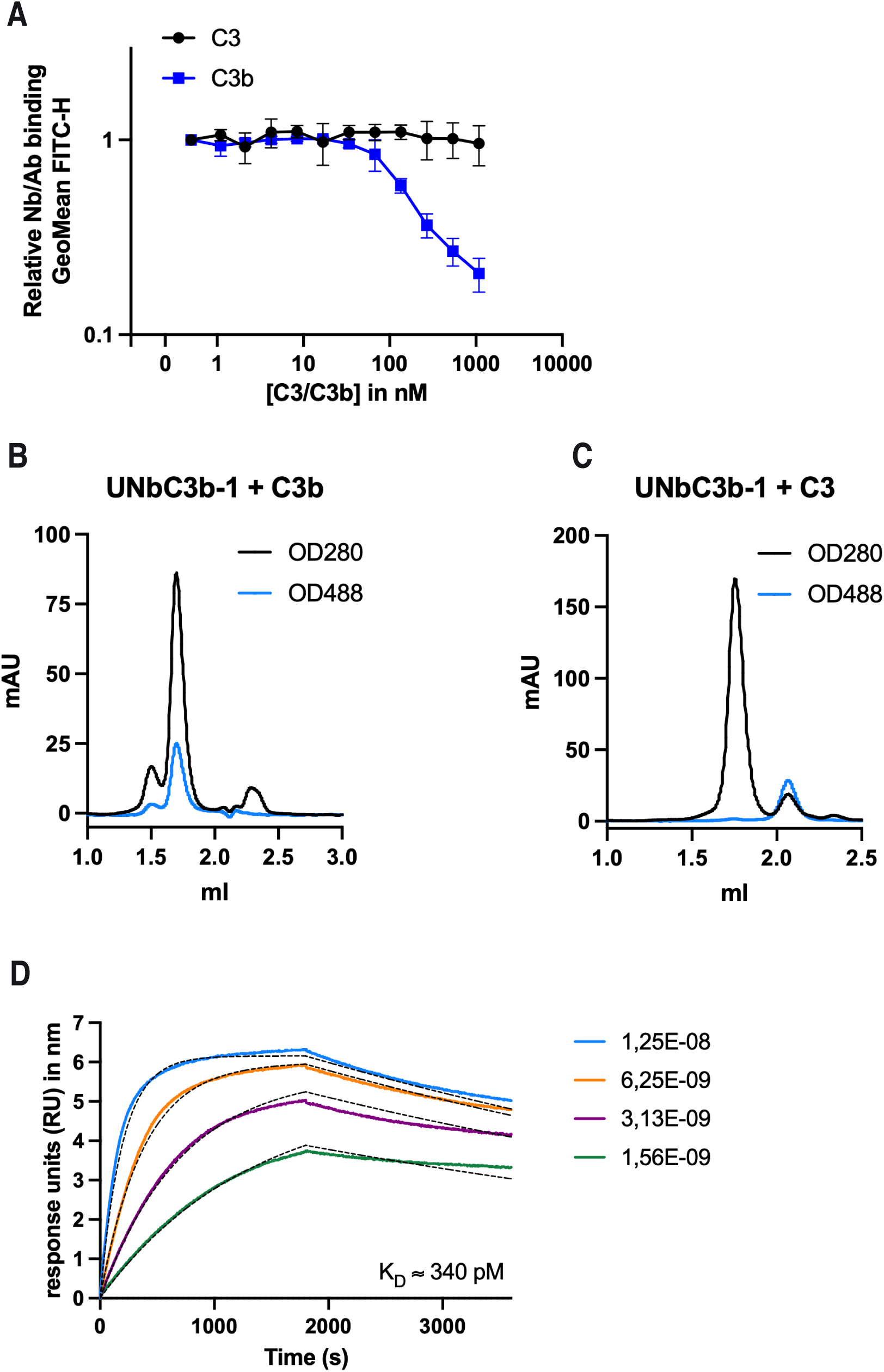
UNbC3b-1 specifically binds C3b with high affinity. (A) Binding of 100 nM UNbC3b-1-a488 to C3b-beads in the presence of increasing concentrations of C3b and C3 in solution, as detected by flow cytometry. Data represent mean ± SD of three individual experiments. (B-C) Chromatograms obtained with SEC, using a Superose 6 Increase column and measuring absorbance at an OD of 280 nm (indicating the presence of protein, black) and of 488 nm (indicating the presence of fluorescence, blue). UNbC3b-1-a488 was incubated with C3b (B) or C3 (C). Data represents two individual experiments. (D) Binding affinity curves of UNbC3b-1 obtained with a White Fox optic surface plasmon resonance (FO-SPR) sensor, using streptavidin FO-SPR probes, UNbC3b-1-biotin (330 nM) as an analyte and C3b (1.56 – 12.5 nM) as a ligand. Immobilization of UNbC3b-1-biotin was performed for 300 seconds, association was measured for 1800 seconds and dissociation for 7200 seconds, all at 26°C, while shaking at 1000 rpm. Experimental data is shown in colors and model fit in black dotted lines. Curves were fitted with TraceDrawer, using a 1:1 model with global B_max_ (6.32), global k_on_ (4.03×10^5^ M^-1^s^-1^), global k_off_ (1.37×10^-4^ s^-1^) and a constant BI (0.0 Signal). Data are representative for two individual experiments.

To facilitate further characterization of UNbC3b-1 and allow direct detection and coupling of this nanobody, we cloned the UNbC3b-1 gene in a vector that replaced the Myc-6×His-tag for an LPETG-6×His-tag. UNbC3b-1 could now be used directly (UNbC3b-1), or when labeled after sortagging with biotin (UNbC3b-1-bio) or fluorophore Alexa Fluor 488 (UNbC3b-1-a488) (21). We confirmed that a change in expression system and the introduction of these tags, did not affect the ability of UNbC3b-1 to bind C3b (**SFig. 3**).

To assess C3b-specificity with a different set-up, we performed size exclusion chromatography (SEC). As controls C3b, C3 and UNbC3b-1-a488 were applied to the column alone. The obtained chromatograms indicated that C3 and C3b elute in fractions between 1.5-2.0 mL, while UNbC3b-1 elutes in fractions between 2.0-2.2 mL (**SFig. 4A-C**). When UNbC3b-1-a488 was incubated with C3b in a 1:1 molar ratio, UNbC3b-1 eluted together with C3b in the fractions between 1.5-2.0 ml, indicating complex formation in solution (**Fig. 3B**). Complex formation was not observed when UNbC3b-1 was incubated with C3 (**Fig. 3C**).

**Figure 4:**
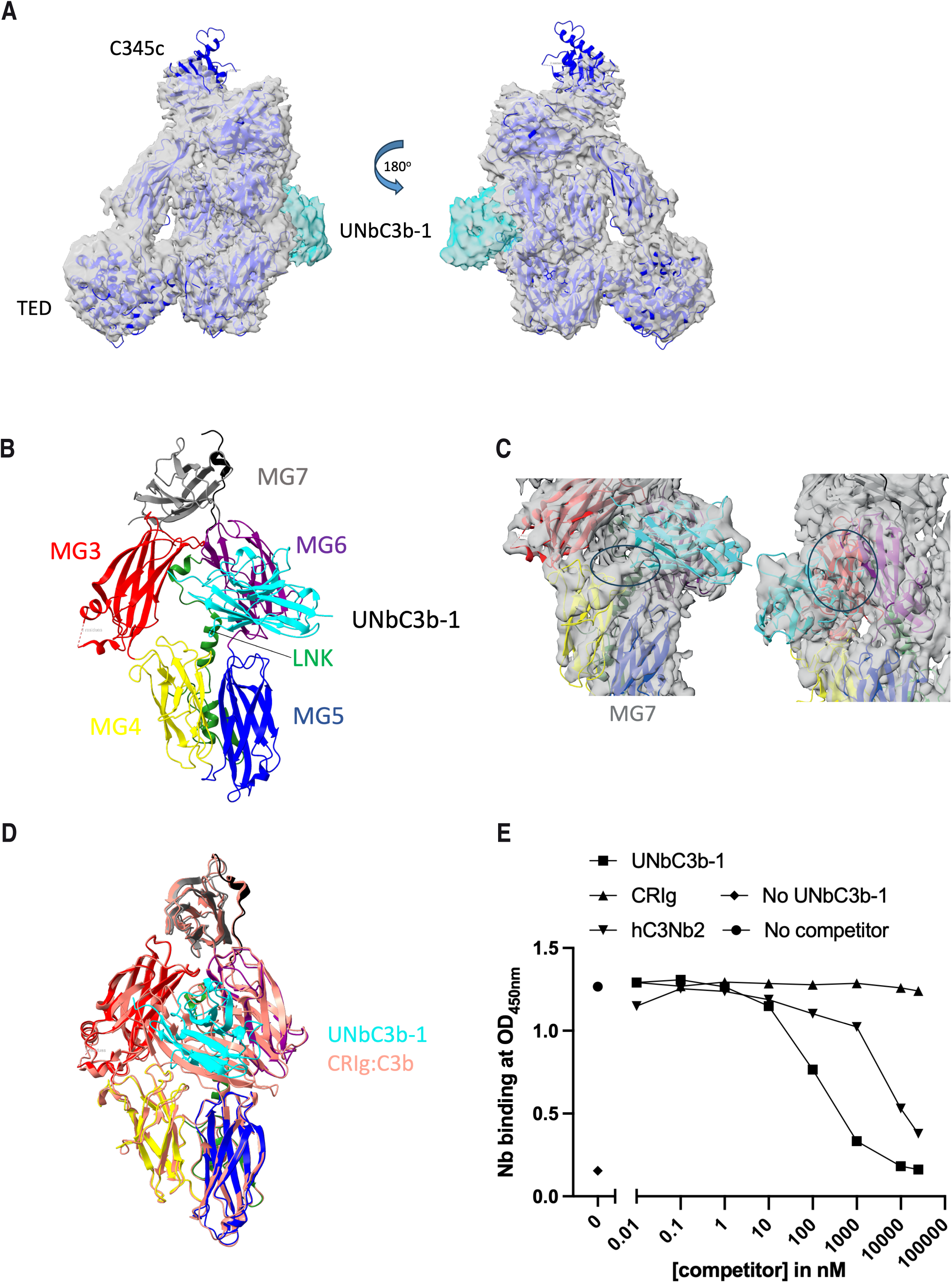
UNbC3b-1 binds the substrate binding face of C3b at the MG3,4,6 interface. (A) The crystral structure of C3b (PDB 2i07) docked into the cryo-EM generated density map. Only one portion of the map was unassigned to C3b (green) and corresponded well to nanobody UNbC3b-1. (B) Rigid body fitting of C3b and UNbC3b-1 resulted in a molecular model of the binding interface, which located to the C3b MG3-6 ring. (C) Closer inspection of the binding interface showed direct connections in the density map. However, resolution was not sufficient to determine the exact amino acid interactions. (D) A comparison of the UNbC3b-1:C3b model with the structure of CRIg bound to C3b showing that the binding regions overlap without major changes to the MG3-6 ring. (E) Competition ELISA with C3b coated on microtiter plates that were incubated with 10 nM UNbC3b-1-biotin and different concentrations of CRig and hC3Nb2. UNbC3b-1-biotin binding was measured using streptavidin HRP. Samples without a competitor and without UNb-C3b-1 were taken along as controls. Data is a representative figure for two individual experiments.

We next assessed the binding kinetics of UNbC3b-1 to C3b by using surface plasmon resonance (SPR). Here, we captured UNbC3b-1-bio onto streptavidin-coupled probes and incubated those with C3b and buffer to measure association and dissociation rates. Fitting this with a 1:1 Langmuir model results in an on-rate (k_on_) of ∼4×10^5^ M^-1^s^-1^, an off-rate (k_off_) of ∼1.3×10^-4^ s^-1^, and a calculated equilibrium binding affinity constant (K_D_) of ∼340 pM. However, as the obtained curves did not completely fit with 1:1 model, the affinity obtained from this experiment is merely a rough indication of its actual affinity (**Fig. 3D**). Taken together, we confirm that UNbC3b-1 is a high affinity, C3b specific nanobody, that binds to C3b in solution and on a surface.

### UNbC3b-1 binds the interface of MG3, 4, & 6 on the C3b substrate binding surface

To determine where UNbC3b-1 binds to C3b, we used cryogenic electron microscopy (cryo-EM) and single particle analysis (SPA) to determine the structure of the UNbC3b-1:C3b complex. We combined UNbC3b-1 in a 3:1 molar ratio with C3b and prepared it for imaging by plunge freezing. Initial screening showed the complex embedded in vitreous ice with a sufficient particle density for data collection (**Fig. S5A**). Image collection resulted in 1568 micrographs that were then computationally processed using the CryoSPARC software suite (22). The built-in blob picker was used to select particles for template generation. Several rounds of cleaning via 2D classification resulted in 253,744 particles with 2D classes containing C3b-like features (**Fig. S5B**). Five *ab-initio* models were generated, revealing one class of 76,815 particles that had structural features similar to C3b (PDB 2I07) that also included an additional unknown density (**Fig. S5C**). Further cleaning of this set of particles using homogeneous refinement and non-uniform refinement resulted in a final density map with a global resolution of 4.2 Å (**Fig. 4A; Fig. S5D**). This map allowed the clear determination of the binding region of the UNbC3b-1 nanobody to C3b. Using rigid body fitting, we were able to dock the crystal structure of C3b (PDB 2I07) and an AF2-predicted structure of UNbC3b-1 into the density map (**Fig. 4A**). We show that the nanobody locates to the interface of the MG3, MG4, and MG6 domains of C3b (**Fig. 4B**), likely with direct interaction with MG3 and MG6 (**Fig. 4C**).

To explain UNbC3b-1’s specificity to C3b, an initial comparison of published C3 and C3b structures showed that there are limited changes in MG3, MG4, and MG6 domains, and thus no obvious reason for the C3b specificity. A query of published structures of C3/C3b binders resulted in a number of proteins that bind in the vicinity of UNbC3b-1: nanobodies hC3Nb1-3 and CRIg (23–26). Interestingly, only two of these binders overlaps with the binding region of UNbC3b-1. hC3Nb2 interacts with only the MG3 and MG4 domains of C3b, while UNbC3b-1 also interacts with MG6. CRIg overlaps significantly with UNbC3b-1 density, and its specificity to C3b requires MG3 and the LNK domain, which would be possible for UNbC3b-1 **(Fig. 4D)**. Unfortunately, we could not clearly resolve this interaction in our density map, thus limiting our interpretation. Experimentally, we used a competition ELISA to see if hC3Nb2 or CRIg can compete with UNbC3b-1. hC3Nb2 partially competes with the binding of UNbC3b-1 to C3b, which is supported by the partial overlap in binding region. However, CRIg did not compete with UNbC3b-1 in this assay, possibly due to the difference in affinity for the epitope (**Fig 4E**, low µM for CRIg vs. low nM for UNbC3b-1).

In conclusion, we show that UNbC3b-1 binds to the C3b beta subunit at the interface of MG3, MG4, and MG6, and that this region directly overlaps with the CRIg binding domain.

### UNbC3b-1 inhibits the AP by blocking convertase function

Since CRIg has been shown to block the AP at the level of the convertase (23), we hypothesized that UNbC3b-1 would also inhibit the AP. Thus, we first determined if UNbC3b-1 could inhibit complement activity using erythrocyte lysis assays for the AP and CP. Indeed, UNbC3b-1 inhibited the AP in a dose-dependent manner but did not inhibit the CP **(Fig. 5A & B)**. Using the same assay, we confirmed that the fluorescently-labeled versions of UNbC3b-1 also inhibited the AP **(SFig. 6A)**. To unravel the mechanism by which UNbC3b-1 blocks the AP, we next performed complement deposition ELISAs to determine at what level the nanobody inhibits complement activity. Here we observed that UNbC3b-1 prevents AP-mediated C3b deposition (**Fig. 5C)** and not CP-mediated C3b deposition (**Fig. 5D**). Likewise, UNbC3b-1 prevents C5b-9 deposition in the AP, but not in the CP (**SFig. 6B**).

**Figure 5:**
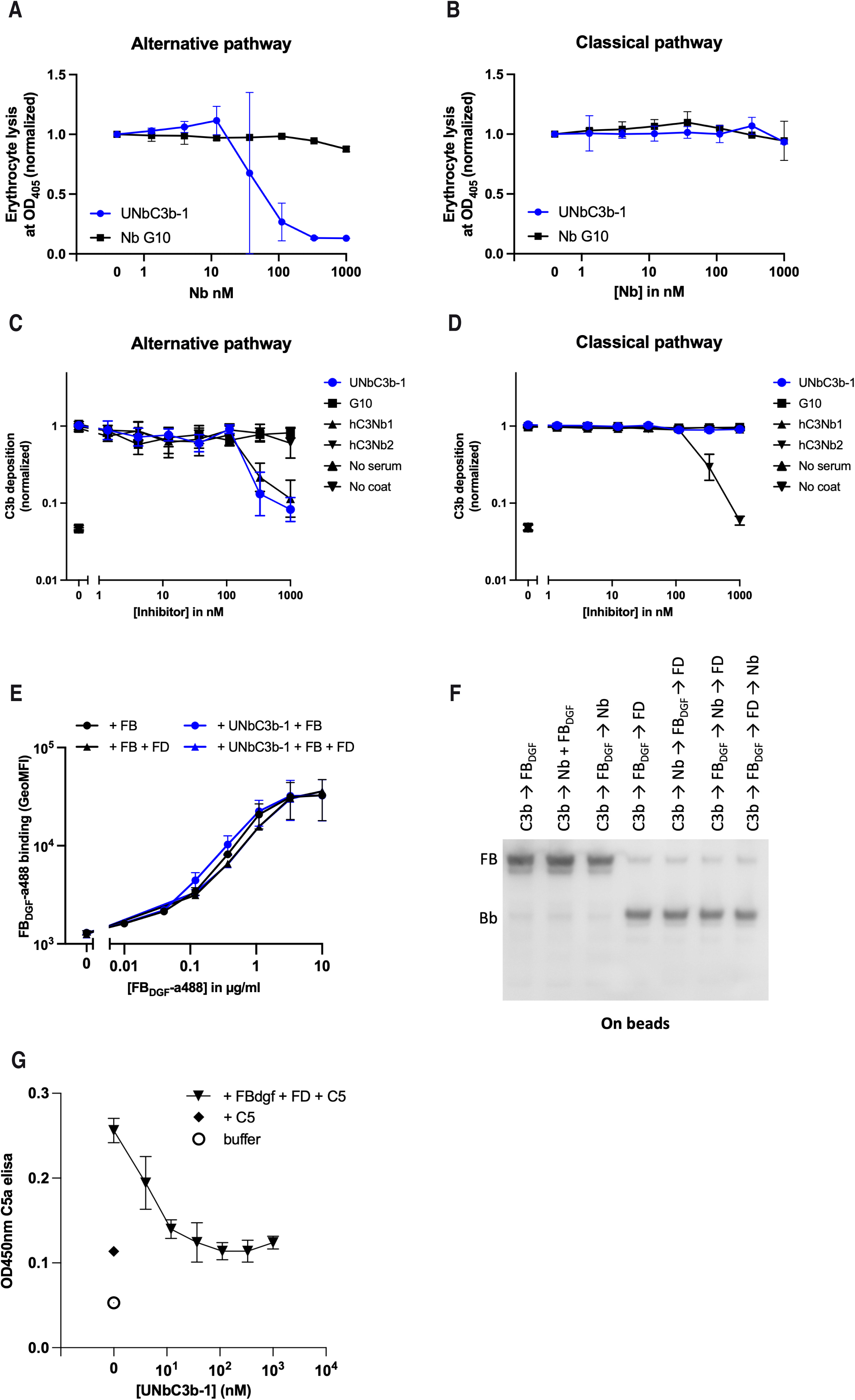
UNbC3b-1 inhibits the AP by blocking convertase functioning. (A) AP-mediated hemolysis of 2% rabbit erythrocytes incubated with 10% human serum and nanobodies UNbC3b-1 and G10 in different concentrations (B) CP-mediated hemolysis of 2% antibody-opsonized sheep erythrocytes incubated with 2.5% human serum and nanobodies UNbC3b-1 and G10 in different concentrations. The OD450 of the supernatant was detected as a measure of erythrocyte lysis. (C) Alternative pathway complement ELISA in which an LPS-coated microtiter plate was incubated with 10% human serum and different concentrations of inhibitors. (D) Classical pathway complement ELISA in which an IgM-coated microtiter plate were incubated with 2.5% human serum and different concentrations of inhibitors (C-D) Deposition of complement activation product C3b on the plate was measured using the WM-1 DIG labeled anti-C3 antibody and an anti-DIG-PO secondary antibody. (A-D) Data points were normalized to the condition without inhibitor and graphs represent mean ± SD of three individual experiments. (E) Interaction of fluorescently labeled FB_DGF_ with C3b beads in the presence of UNbC3b-1 and/or FD. Binding of FB_DGF_-a488 to the beads was assessed using flow cytometry. Data represent mean ± SD of two individual experiments. (F) Cleavage of FB by FD in the presence or absence of UNbC3b-1, using C3b-beads. Proteins were added in different orders (see lanes) with 10 minutes incubation steps in between, indicated with an arrow. Bb formation was assessed by western blot using polyclonal anti-FB-HRP labeled antibodies. Bb can be distinguished from uncleaved FB based on the height of the band (corresponding to protein size). Figure represents data of two individual experiments. (G) C5a ELISA to measure C5a formation using C3b-beads and convertase components. Components were sequentially added to C3b-beads with 10-minute-incubation steps at RT in between. Components were added in the following concentrations 20 µg/mL FB_dgf_, 5 µg/mL FD, 4.1 – 1000 nM UNbC3b-1 and 20 µg/mL C5. C5a levels were measured by adding six times diluted supernatant to a C5a ELISA kit. Data represent mean ± SD of two individual experiments.

**Figure 6:**
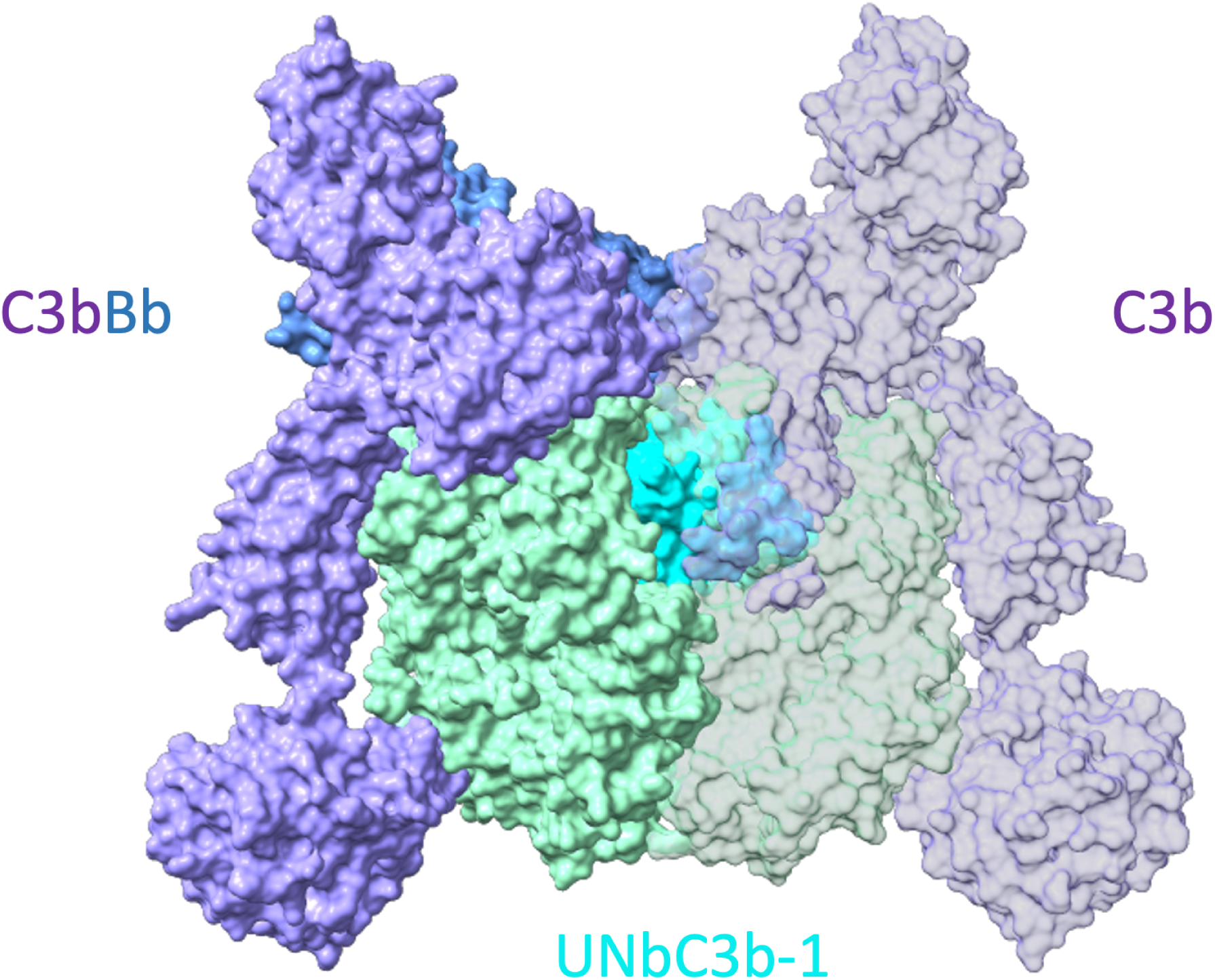
UNbC3b-1:C3b binds the substrate binding surface of the AP C3bBb convertase. A comparison of the UNbC3b-1:C3b structure with the SCIN-stabilized AP convertase (2WIN) showing that UNbC3b-1 locates to the C3b:C3b binding interface, likely interfering with substrate binding.

We then assessed if UNbC3b-1 inhibits complement activity by preventing the formation of the AP C3 convertase. These convertases are formed when covalently deposited C3b molecules are bound by FB (proconvertase, C3bB) and processed by FD (active convertase, C3bBb). First, we studied whether UNbC3b-1 could prevent formation of proconvertases. To do so, C3b-beads were incubated with UNbC3b-1 and a fluorescently labeled double gain of function mutant of FB (FB_DGF_) that binds C3b more stable than WT FB. This is required to prevent the detachment of FB in washing steps. FB_DGF_ binding to C3b-beads was assessed using flow cytometry. Interestingly, UNbC3b-1 did not inhibit C3bB formation since FB_DGF_ binding to C3b was unaffected in the presence of the nanobody (**Fig. 5E**). Also, when FD was added to generate active convertases from proconvertases, UNbC3b-1 did not block Bb binding to the C3b-beads (**Fig. 5E**). To test if FD could still cleave FB into Ba and Bb in the presence of UNbC3b-1, C3b-coupled beads were incubated with FB_DGF_, FD and UNbC3b-1, in varying orders. In all tested conditions in which FD was present, we observed that FB was cleaved into Bb (**Fig. 5F**). The presence of UNbC3b-1 did not influence this, indicating that UNbC3b-1 does not prevent formation of an active convertase enzyme.

Finally, we assessed if UNbC3b-1 inhibits substrate cleavage by the convertase. To do so, we incubated C3b-beads with UNbC3b-1, FB_dgf,_ FD and C5, and subsequently measured the presence of C5a in the supernatant using a sandwich ELISA. Indeed, we observed that UNbC3b-1 inhibits C5 cleavage by C3bBb-beads (**Fig. 5G**). Taken together, UNbC3b-1 is an AP specific inhibitor that blocks C3 and C5 cleavage by AP convertases.

## DISCUSSION

With the identification of UNbC3b-1 as a C3b-specific nanobody, this study adds a novel complement-targeting molecule to the field of complement research. Conversion of C3 into C3b is an essential step in complement activation. Therefore, molecules that can distinguish the activated C3b product from precursor C3 are very helpful to detect and further understand the molecular mechanisms at play during complement activation.

Next to UNbC3b-1, three other nanobodies that bind C3b (non-exclusively) have been described, hC3Nb1 (25), hC3Nb2 (26), and hC3Nb3 (24). Like UNbC3b-1, hC3Nb1 inhibits the AP but by a different mechanism. hC3Nb1 binds to C3b MG7 and prevents both the formation of the proconvertase (C3bB) and binding of the substrate C3 to the convertase. hC3Nb3 is unique because it binds to the C345c domain of C3/C3b/C3c and inhibits only the AP C3 convertase but also the AP and CP C5 convertase. In this study, we show that hC3Nb2 has a partially overlapping epitope with UNbC3b-1 and it can partially compete with UNbC3b-1 in a competition ELISA. However, while UNbC3b-1 acts solely on the AP, hC3Nb2 binds C3/C3b/C3c and blocks all three complement pathways by preventing the association of C3 with the convertase. It also explains why hC3Nb2 differs from UNbC3b-1 based on pathway specificity. hC3Nb2 recognizes C3 and C3b by binding to the MG3 and MG4 domains while UNbC3b-1 specifically inhibits the AP by selectively binding to C3b at an interface that is more centered in the MG3-6 ring, preventing the association of C3 with the AP C3 convertase.

Specific binding to C3b has also been described for monoclonal antibodies S77, 3E7, H17, anti-C3-9, and 4C2 that all specifically bind C3b (and not C3) and inhibit the alternative pathway (27–30). Despite the overlap in target and pathway specificity, the mechanism of inhibition of UNbC3b-1 differs from the previously described monoclonal antibodies, since all these block binding of FB to C3b, while we show that UNbC3b-1 does not interfere with FB binding.

Interestingly, a comparison of our density map and UNbC3b-1:C3b complex with known binders of C3b identified that the complement receptor CRIg has an overlapping binding region (PDB 2ICF, **Fig. 4D**). CRIg is found mainly on macrophages in the liver that respond to opsonized bacteria via C3b and iC3b, and it has been shown to inhibit the AP (23). Future studies could explore if UNbC3b-1 can block the interaction of C3b-opsonized particles with CRIg. Alternatively, UNbC3b-1 has the potential to compete with CRIg, providing a new tool that could dampen the macrophage phagocytic response. UNbC3b-1 specificity is likely similar to that of CRIg, which is due to epitopes in C3b exposed by movement of the MG3 and LNK domains caused by the major rearrangement that occurs following the cleavage of C3a.

Based on our model, we have shown that UNbC3b-1 acts on the AP convertase. A comparison of the SCIN-stabilized AP C3 convertase (PDB 2WIN;(31)) with our model shows that UNbC3b-1 binds at the substrate binding interface, preventing the association of C3 with the convertase (**Fig. 6)**. In addition, due to the high affinity of UNbC3b-1, it is likely that all C3b molecules are bound to the nanobody and thus prevents the activity of the AP convertase. However, the structure of the AP C5 convertase is unknown, and thus we do not know the confirmation or binding site(s) of the additional C3b molecules necessary for substrate conversion, and thus the full impact of UNbC3b-1 on the AP C5 convertase remains a question.

With the identification of nanobody UNbC3b-1, this study adds a novel C3b binding molecule to the field that is distinct from previously described anti-C3b antibodies. The increasing number of C3- and C3b-binding molecules with different mechanisms of inhibition and pathway specificity, aids in unravelling the complex molecular mechanisms at play in C3 and C5 convertase formation and functioning. Finally, since UNbC3b-1 efficiently inhibits the AP of the complement system, this molecule might generate new insights for the development of novel therapeutic complement inhibitors.

## EXPERIMENTAL PROCEDURES

### Proteins, serum and bacterial strains

Normal human serum from ±20 healthy donors was pooled, as previously described (32). *E. coli* strains MG1655, BL21, Rosetta 2 (DE3), BL21 (Merck) and TG1 (Agilent) were used. Complement components and inhibitors were obtained from sources listed in Table II. Complement component Bb was produced by incubating C3b-PEG11-biotin with 6×His-tagged FB and 6×His-tagged FD for 1 hour at 37°C, in a molar ratio of 1:10:3. Next streptavidin-6×His was added in a 1.75:1 molar ratio compared to C3b-PEG11-biotin for 1 hour at 4°C to couple C3b-PEG11-biotin with a 6×His-tag. Next, the sample was applied to a 1 mL HisTrap FF column (GE Healthcare) and the flowthrough containing Bb was captured, dialyzed to phosphate buffered saline (PBS) and stored at −80°C.

**Table II:** sources of complement proteins and inhibitors.

| # | Component | Source | Company/in-house | references |
| --- | --- | --- | --- | --- |
| 1 | C3 | Human plasma | In-house | (10) |
| 2 | C3b | Human plasma | In-house | (47) |
| 3 | C3b-PEG11-biotin | Human plasma | In-house | (9) |
| 4 | C3b-PEG4-DBCO | Human plasma | In-house | (33) |
| 6 | C3a |  | Complement Technology |  |
| 7 | C3a desArg |  | Calbiochem |  |
| 8 | iC3b-PEG11-biotin | Human plasma | In-house | unpublished |
| 9 | C3c |  | Complement Technology |  |
| 10 | C3dg | Recombinant | In-house | unpublished |
| 11 | C3d |  | Complement Technology |  |
| 12 | FB<sub>WT</sub> | Recombinant | In-house | (9) |
| 13 | FB<sub>WT</sub>-a488 | Recombinant | In-house | unpublished |
| 14 | FB<sub>DGF</sub> |  | In-house, plasmid kind gift from Piet Gros | (48) |
| 15 | FB<sub>DGF</sub>-a488 | Recombinant | In-house | unpublished |
| 16 | FD | Recombinant | In-house | (9) |
| 17 | Bb | Recombinant | In-house | See methods |
| 18 | C4 |  | Complement Technology |  |
| 19 | C4b |  | Complement Technology |  |
| 20 | C5 | Recombinant | In-house | (38) |
| 21 | C5a |  | Bachem, Zwitterland |  |
| 22 | C5b6 |  | Complement Technology |  |
| 23 | C6 | Recombinant | In-house | (49) |
| 24 | hC3Nb1 | Recombinant | In-house, based on | (25) |
| 25 | hC3Nb2 | Recombinant | In-house, based on | (26) |
| 26 | hC3Nb3 | Recombinant | In-house, based on | (24) |
| 27 | CRig |  | Kindly provided by Genentech (south San Fransico, CA, USA) | (9) |
| 28 | Eculizumab | Recombinant | In-house | (38) |

### C3b-opsonized bacteria for immunizations

*E. coli* bacteria strains MG1655 and BL21 were covered wth human C3b molecules, similar as described before (33). To this end, C3b was site specifically labeled with a maleimide linker (10) containing a dibenzocyclooctyne (DBCO) group at its thioester domain. To prepare the bacterial strains, a single colony from *E. coli* MG1655 and BL21 was grown in Luria-Bertani (LB) medium to stationary phase after which a subculture of 1/100 was grown overnight (O/N) in LB medium containing 2 mM azide-modified keto-deoxy-octulosonate (KDO-N_3_), to allow metabolic incorporation of KDO-N in the LPS of these strains. Next, bacteria were washed twice with PBS to remove unincorporated KDO-N_3_, resuspended in PBS containing 17% glycerol and frozen at −20°C. Frozen bacteria were treated with γ-irradiation at a dose of 10.3-10.8 kGy for ±1.5 h (Synergy health Ede B.V. a Steris company), to ensure that bacteria incorporated with KDO-N_3_ were no longer viable. To couple with C3b molecules, γ-irradiated bacteria were thawed, washed, and 2×10 bacteria/ml were incubated O/N, at 4°C with 2.66 µM C3b-PEG4-DBCO. The DBCO in C3b-PEG4-DBCO covalently couples with the azide (N_3_) from KDO-N_3_, resulting in the coupling of C3b molecules to the bacterial membrane (KDO-C3b). The next day, KDO-C3b bacteria were washed 3× with PBS to remove unbound C3b-PEG4-DBCO. To obtain bacteria with C3b deposition in high densities, KDO-C3b bacteria were diluted 5× in Hepes buffer and three rounds of natural C3b amplification were performed. For each round we added FB (5 µg/ml), FD (0.5 µg/ml), and uncleaved C3 (100 µg/ml) and incubated the bacteria for 15 minutes at 37°C. After each round bacteria were spun down, supernatants were removed and new components were added. After the third round of amplification bacteria were washed 3× with Hepes buffer and subsequently split in three equal parts. Next, bacteria were crosslinked with bis(sulfosuccinimidyl)suberate (BS3) (Thermo Scientific), paraformaldehyde (PFA) or not crosslinked at all, to present deposited C3b in different ways to the llamas. Crosslinking with BS3 (5 mM) was performed for 90 minutes at room temperature (RT), the reaction was quenched with 50 mM Tris, for 15 minutes at RT, after which bacteria were washed and resuspended in hepes buffer. Crosslinking with PFA (1%) was performed for 15 minutes at RT, bacteria were washed and resuspended in hepes buffer.

### Immunizations

Two llamas (Lama glama), named Ines and Lotte, were immunized by administration of human C3b on day 0, 14, 28, 35, 57, and 71 by a trained veterinarian at Preclinics GmbH (Potsdam, Germany). All procedures were according to European animal welfare laws and regulations and al animals remained alive after immunization and blood collection. The first four injections each contained 12.5 µg purified C3b in solution. The last two injections contained *E. coli* bacteria opsonized with human C3b. Per injection, both llamas received ±1.5×10^9^ γ-irradiated *E. coli* bacteria opsonized with a high density of C3b molecules. The llama named Ines received *E. coli* strain MG1655 and the llama named Lotte received *E. coli* strain BL21. The samples contained a mixture of the different crosslinking methods. All injections were combined with FAMA adjuvant (Gebru Biotechnic GmbH). To obtain llama serum and PBMCs, blood was drawn on day 0, day 43 and day 99.

### Phage library construction

A nanobody phage display library was generated for each llama. mRNA was isolated form PBMCs (day 43 and day 99), and cDNA was created using the SuperScript™ IV First-Strand Synthesis System kit (Thermo Scientific, 18091050). We amplified the cDNA encoding for the heavy-chain only antibodies by PCR, using primers annealing to framework 1 (FR1) and constant heavy chain 2 (CH2) domains (34). Next, DNA fragments encoding for the nanobody genes were digested and ligated into phagemid pPQ81 vector (derived from pHEN1 (35)). This fused the nanobodies to a Myc- and 6×His-tag and to pIII coat protein of M13 bacteriophage. Next, phage display libraries were transformed into competent *E. coli* TG1 bacteria (Agilent), by electroporation with a MicroPulser and corresponding electroporation cuvettes (Bio-Rad). To estimate library sizes, a serial dilution of transformed bacteria was spotted on LB agar plates and colony forming units per mL (CFU/ml) were counted.

### Phage production and purification

Phages were produced according to standardized protocols (36), using the original phage display libraries obtained after immunization (day 43 and day 99) as starting material for round 1, and the bacterial libraries obtained after the first round of phage display as starting material for round 2. Briefly, *E. coli* TG1 libraries were grown in 2× concentrated yeast extract tryptone (2YT) medium, supplemented with 2% glucose and 100 µg/mL ampicillin. Next, kanamycin resistant VCSM13 helper phage (Agilent) was added and phages were purified via two rounds of polyethylene glycol precipitation.

### Phage display panning

We performed two rounds of phage display, following standardized procedures (37), with in total 9 different selection strategies (**Table I**), combining different methods of antigen presentation and negative selections. Selection strategy #6 yielded nanobody UNbC3b-1. Wells presenting C3b (see details below) were blocked with 200 µL/well 4% skimmed milk (Marvel) in PBS, for 1 hour at RT, while shaking. Next, wells were incubated with phages at concentrations of ≍10^10^ CFU/well (round 1) and ≍10^9^ CFU/well (round 2) in 2% Marvel in PBS, for 2 hours, while shaking. In case of negative selections with purified proteins (e.g. selection strategy #1), phages were preincubated for 30 minutes at RT on a spinning wheel with C3 (30 nM), C5 (30 nM) and/or C3b (30 nM), before being added for 2 hours to the C3b presenting wells. In case of negative selections with beads (e.g. selection strategy #7), handling was the same except that low density C3b-beads (5.0 µL/sample, at 10 mg/mL), including all phages bound by them, were removed from the sample using a magnet, prior to adding them to C3b presenting wells. Next, wells were washed 20× with 200 µl PBS + 0.05% tween-20 (PBS-T), with two 10-minute incubations (shaking) in-between washing steps. Finally, wells were washed 3 additional times using PBS. Next, phages bound to C3b were eluted using 100 µL triethanolamine solution (0.1 M, pH>10), for 15 minutes at RT, while shaking and elutions were neutralized using 50 µL Tris/HCl (1 M, pH=7.5). Phages were rescued by infecting them in *E. coli* strain TG1 bacteria at OD_600_ of 0.5, for 30 minutes, at 37°C, without shaking. Next, bacteria were grown O/N at 37°C, while shaking, in 2YT medium, supplemented with 2% glucose and 100 µg/mL ampicillin. The next day, bacterial libraries containing the genetic material of C3b-binding nanobodies were stored in 20% glycerol at −80°C. To estimate the number of phages used as starting material (input) and the number of phages in the eluate (output), phages were serially diluted, added to *E. coli* TG1 bacteria to allow infection and spotted on LB agar plates supplemented with 2% glucose and 100 µg/mL ampicillin. After an O/N incubation at 37°C, CFU/mL were counted and estimated library sizes were calculated.

### C3b presentation during phage display

When presenting C3b on plates (e.g. selection strategy #1, Table I), Nunc Maxisorp plates (VWR 735-0083, Thermo Fisher Scientific) were coated O/N, at 4°C, without shaking, with 100 µL/well C3b at a concentration of 0.1, 1.0 or 5.0 µg/mL. When presenting C3b in high densities on beads, magnetic streptavidin-coated beads (0.1, 0.2, 0.8, 2.0 or 4.0 µL/sample, at 10 mg/mL, Dynabeads M-270 Streptavidin, Invitrogen) were diluted 50× and coupled O/N at 4°C with 1 µg/mL C3b-PEG11-biotin (C3b-bio) in veronal buffered saline + 145 nM NaCl, pH 7.4 (VBS), supplemented with 2.5 mM MgCl_2_ and 0.05% Tween-20 (VBS/MgCl_2_/Tween). After O/N coupling, unbound C3b-bio was washed away. To obtain beads with low densities of C3b, used for counter selections, protocols were the same, except beads were diluted 3.125× instead of 50× before adding C3b-bio.

### Periplasmic nanobody production for screening

Bacterial output libraries (*E. coli* TG1) were plated on LB agar plates supplemented with 2% glucose and 100 µg/mL ampicillin, incubated O/N at 37°C, to obtain single colonies. Next, single colonies were randomly picked and 92 wells containing 90 µL 2YT medium, supplemented with 2% glucose and 100 µg/mL ampicillin were inoculated with one colony/well. As negative controls, two wells were not inoculated (EM), and two wells were inoculated with a bacterium expressing a nanobody with an irrelevant target (IRR) or a bacterium containing an empty vector (PER). Bacteria were grown O/N at 37°C, without shaking. The next day, subcultures (1/100) were made in 900 µL 2YT medium with 0.1% glucose and 100 µg/mL ampicillin and grown until a clear growth difference was observed between negative controls (EM) and other wells (approximately 2.5-3 h). Then, bacterial cultures were induced by adding 1 mM isopropylthio-β-galactoside (IPTG) and grown O/N, at RT, while shaking. The next day, cultures were spun down for 10 minutes at 6000 g, and pellets were resuspended in 120 µL PBS. Next, bacterial suspensions were frozen at least O/N at −20°C or >30 minutes at −80°C, before thawing. After thawing, samples were spun down 2× for 10 minutes at 6000 g and supernatants, containing the periplasmic extracts with nanobodies in unknown concentrations, were collected and stored at −20°C, until further use.

### Nanobody productions

Nanobodies containing a Myc- and 6×His-tag (further referred to as MH-clones) were produced in the periplasm of *E. coli* (38). For this, *E. coli* Rosetta 2 (DE3) BL21 cells (Merck) were transformed with pPQ81 vectors encoding the nanobody genes and C-terminal Myc- and 6×His-tags. Next, bacteria were grown O/N, at 37°C, while shaking, in 2YT medium, supplemented with 2% glucose and 100 µg/mL ampicillin. The following day, subcultures (1/20) were grown until they reached an OD_600_ of 0.6-0.9, in 2YT medium supplemented with 0.1% glucose and 100 µg/mL ampicillin. Nanobody expression was induced for four hours at 37°C, while shaking, by adding 1 mM IPTG. Next, cultures were spun down for 10 minutes at 6000 g, pellets were dissolved in PBS and frozen for >30 minutes at −80°C, or O/N at −20°C. Frozen pellets were thawed to release periplasmic extracts in the supernatant, which were collected after 15 minutes of centrifugation at 6000 g. Nanobody UNbC3b-1 (clone B5) containing an LPETG-6×His-tag (further referred to as UNbC3b-1) was cloned into a pcDNA 3.4 TOPO™ vector (Thermo Fisher Scientific) and expressed in EXPI293F cells (Thermo Fisher Scientific), similar as described for complement protein C5 in Struijf et al. (38), except that here we added 1 µg/ml DNA/ml cells for transfection.

### Nanobody purifications

MH-clones were isolated using immobilized metal-affinity chromatography. For this, supernatants containing the nanobodies were incubated with ROTI Garose-His/Co beads (Roth) for 30 minutes, at RT, on a spinning wheel, to bind nanobodies via their 6×His-tag. After three washing steps with PBS-T, nanobodies were eluted with 150 mM imidazole. For purification of UNbC3b-1, EXPI293F supernatant was buffer exchanged to 50 mM Tris, 500 mM NaCl, pH8.0 via dialysis and 30 mM imidazole was added before application to a HiTrap Chelating column (GE Healthcare). The column was washed with 30 mM imidazole, before UNbC3b-1 was eluted with a 30 to 250 mM imidazole gradient, using the Äkta Pure protein chromatography system (GE Healthcare). Finally, all nanobodies were dialyzed to PBS, concentrations were measured at A_280_ using a Nanodrop One (Thermo Fisher Scientific) and nanobody size, purity and concentrations were verified using SDS-PAGE, stained with InstantBlue Safe Coomassie stain (Sigma Aldrich).

### Nanobody labeling with biotin and fluorophores

We labeled UNbC3b-1 with biotin (further referred to as UNbC3b-1-bio) and fluorophore Alexa488 (further referred to as UNbC3b-1-a488) via sortagging (21). First, UNbC3b-1 was dialyzed against 50 mM Tris, 150 mM NaCl, pH8.0 (sortase buffer). Next, UNbC3b-1 was incubated with His-tagged Sortase A7 (His-TEVG-SrtA7, (21)) and GGG-azide (Genscript) in a 7:1:70 molar ratio, in sortase buffer supplemented with 10 mM CaCl_2_, for 2 hours, at 4°C, while shaking. 20mM imidazole was then added and samples were applied to a HIS-Trap FF column (GE Healthcare), to remove UNbC3b-1 that did not get sortagged with GGG-azide, and flow-through, containing UNbC3b-1-GGG-azide was collected. The UNbC3b-1-GGG-azide was subsequently incubated for 2 hours, at RT, while shaking, with a 5× molar excess DBCO-AlexaFluor488 (a488) (JenaBioScience) or DBCO-PEG4-biotin (Santa Cruz Biotechnology) in sortase buffer. Finally, UNbC3b-1-a488 and UNbC3b-1-bio were dialyzed against PBS.

### ELISA: llama antibody and nanobody binding

Nunc Maxisorp plates (VWR 735-0083) were coated O/N at 4°C, without shaking, with 50 µl antigen (C3b, C3, C3a, C4, C4b, C5, C5a, C5b6, C6, Bb, iC3b-biotin, C3c, C3dg, C3d, C3a des-Arg) at a concentration of 2 µg/mL in sterile PBS. The next day, wells were blocked for 1 hour, at RT, with either 200 µL PBS + 4% Marvel (llama immune response) or 80 µL PBS-T + 4% bovine serum albumin (BSA, Sigma) (all others). All following incubations were performed with 50µl sample/well, at RT, for 1 hour, while shaking. Samples were diluted in PBS + 1% Marvell (llama immune response or PBS-T + 1% BSA (all others) and in-between incubations wells were washed 3× with PBS-T. Unless stated here, sample concentrations are indicated on the X-axis in the corresponding figure legend. For the C3b/C3 screening ELISA, periplasmic extracts were diluted 1:5. For the competition ELISA with known C3b-binding molecules, UNbC3b-1-biotin (10 nM) was premixed with competitors in a concentration series. Compounds hC3Nb1, hC3Nb2, hC3Nb3, clone G10 (from this screen) and CRIg were added in 0.1 – 1000 nM; MCP was added in 0.01 – 100 nM. To detect llama antibodies and nanobodies (not via a tag), wells were incubated with 1:2000 QE19 (polyclonal rabbit-anti-nanobody antibodies, QVQ Holding BV), followed by 1:5000 polyclonal donkey-anti-rabbit-PO labeled antibodies (Jackson Immuno research). To detect biotinylated nanobodies, wells were incubated with 1:5000 tetrameric streptavidin coupled to HRP (Southern Biotech). To develop ELISAs substrates were added; 3.7 mM of O-phenylenediamine dihydrochloride + 50 mM Na_2_HPO_4_ 2H_2_O + 25 mM citric acid + 0.03% H_2_O_2_ (llama immune response) or 100 µg/mL tetramethylbenzidine dissolved in DMSO + 100 mM NaOAc + 1.7 mM ureum peroxide (all others). All ELISAs were stopped with 0.5 M sulfuric acid and absorbance (450 nm) was measured with an iMark Microplate Reader (Biorad).

### Sequencing

For sequencing, fresh O/N bacterial cultures were made using 2YT medium, supplemented with 2% glucose and 100 µg/mL ampicillin. In case of sequencing all 96 clones, 5µl of the O/N culture was added to a 96-wells plate filled with LB-agar. Plates were next incubated for 2-3 hours at 37°C, without shaking. Plates were sealed with parafilm and send to Eurofins for Sanger sequencing. In case of sequencing 1-8 clones, bacteria were plated on LB agar plates to obtain single colonies. Next, a single colony was picked and used to inoculate a 5mL culture with 2YT medium, supplemented with 2% glucose and 100 µg/mL ampicillin, that was grown O/N at 37°C, while shaking. The next day, bacterial cultures were spun down and supernatants were removed. Next, plasmids were isolated using the NucleoSpin EasyPure kit (Macherey-Nagel), following instructors’ protocols. DNA concentrations were measured using the MultiSkanGo (Thermo Scientific) and 900-1000 ng DNA was added to 5 µM forward primer, in a total volume of 10 µL milliQ water. Next, samples were sent to Eurofins for sequencing. Using the PipeBio Antibody Sequence Analysis platform resulting nucleic acid sequences were analysed and processed into nanobody amino acid sequences. The nanobody sequences were annotated for frameworks and complementarity determining regions. Sequences were aligned to IgG germlines and clustered using 80% CDR-H3 homology as a cut-off.

### ELISA: complement deposition

Nunc Maxisorp plates (VWR 735-0083) were coated overnight at room temperature, without shaking, with 3 µg/mL human IgM (Millipore) diluted in PBS (classical pathway (CP)) or with 20 µg/mL sonicated LPS from *Salmonella enteritis* (Sigma) diluted in 0.1 M sodium carbonate, pH 9.6 (alternative pathway (AP)). The next day, wells were blocked for 1 hour at RT, while shaking, with 80 µL PBS-T supplemented with 4% BSA. Next, wells were incubated for 1 hour at 37C, while shaking, with 4% (CP) or 40% (AP) human pooled serum, in the presence of 1.3 – 1000 nM UNbC3b-1 and non-inhibitory nanobody G10. Complement inhibitors hC3Nb1 and hC3Nb2 were taken along. C3b deposition was detected using 1:10.000 anti-C3 WM-1 clone digoxigenin (DIG) labeled antibodies (Sigma) and 1:8000 anti-DIG-PO antibodies (Roche). C5b-9 deposition was detected using 1:1000 monoclonal mouse anti-C5b-9 aE11 (produced in our lab, based on (39, 40)) and 1:5000 polyclonal goat-anti-mouse-PO (Southern Biotech). Complement deposition ELISAs were developed similar as described above for the nanobody binding ELISAs.

### Erythrocyte lysis assays

Sheep erythrocytes (shE, Alsever Biotrading) and rabbit erythrocytes (raE, kindly provided by Utrecht University, Faculty of Veterinary medicine) were washed 3x with PBS. For the CP, shE were diluted to a 2% suspension in VBS supplemented with 0.25 mM MgCl_2_ and 0.5 mM CaCl_2_ (VBS++) and shE were opsonized with 1:2000 polyclonal rabbit-anti-sheep IgM antibodies (haemolytic amboceptor (41), shEA). For the AP, raE were diluted to a 2% suspension in VBS, supplemented with 5 mM MgCl_2_ and 10 mM EGTA (VBS/Mg/EGTA). Next, raE and shEA suspensions were incubated for 10 minutes (CP) or 30 minutes (AP), at 37 °C, while shaking, with 2.5% (CP) or 10% (AP) pooled human serum, and different concentrations of nanobodies (1.4 – 1000 nM, MH-clones). As controls, we took along a samples with erythrocytes and buffer (0% lysis), erythrocytes and milliQ water (100% lysis) and samples with serum but no nanobody. Next, samples were spun down for 7 minutes at 3500 rpm. Supernatants were diluted 1:3 with milliQ water and hemoglobin release was measured at and OD of 405 nm, in a flat-bottom plate, using an iMark Microplate Reader (Biorad).

### C3b beads binding assay

Magnetic streptavidin-coupled beads (0.5 µl/sample, at a concentration of 10 mg/mL, Dynabeads M-270 Streptavidin, Invitrogen) were coupled with a high density of C3b molecules, as described above (C3b presentation during phage display). Next, nanobodies (UNbC3b-1-a488 or the eight MH-clones, all at 100 nM) were incubated with the beads and soluble C3 (27 – 270 nM or 0.5 – 1080 nM) or C3b (0.5 – 1080 nM) for 30 minutes at 4°C/RT, while shaking. To detect binding of the eight MH-clones, we next incubated beads with QE19 (QVQ Holding BV) directly labeled with fluorophore Alexa 647 (QE19-a647) for 30 minutes at 4°C, while shaking. In-between incubations, beads were washed 3× with PBS-T to remove excess nanobody, C3, C3b or QE19-a647. Next, beads were fixated for 15 minutes, at RT, by resuspending them in 100µl PBS-T supplemented with 1% PFA. Binding of nanobodies was assessed on a FACS verse flow cytometer (BD).

### FB binding assay

High density C3b beads (1 µL/sample, at 10 mg/mL), prepared as described above, were incubated with 1 µM UNbC3b-1, for 30 minutes at RT, while shaking. Next, beads were washed to remove unbound UNbC3b-1, and incubated with directly labeled FB_DGF_-a488 (0.04 – 10 µg/ml), and FD (0.5 µg/mL), for 30 minutes, at 37°C, while shaking. Next, beads were washed and fluorescence was measured using the FACS Verse flow cytometer (BD).

### FB cleavage assay and western blot

High density C3b beads (4µl beads/sample) were prepared as described above and incubated with FB_DGF_ (100 nM), FD (42.5 nM) and UNbC3b-1 (1 µM). Samples were incubated for 30 minutes, at 37°C, while shaking in VBS/MgCl_2_ buffer. Samples were diluted 1:1 with 2× concentrated reducing SDS sample buffer (0.1 M Tris (pH 6.8), 39% glycerol, 0.6% SDS, and bromophenol blue and incubated for 5 minutes at 95°C. Samples were run on a 4-12% Bis-Tris gradient gel (Invitrogen) for 60 minutes at 200 V. Next, proteins were transferred to 0.2 µM PVDF membranes (Bio-Rat), using the Trans-Blot Turbo transfer system (BioRad). After blotting, membranes were blocked with PBS supplemented with 0.1% Tween-20 and 4% dried skim milk (ELK), for 1 hour, at 37°C, while on a roller bank. Next, blots were incubated with 1:300 diluted polyclonal goat-anti-FB (Complement Technology), washed and incubated with 1:10,000 polyclonal donkey-anti-goat-HRP (Southern Biotech), all in PBS supplemented with 0.1% Tween-20 and 1% ELK. In between all steps, membranes were washed 3× with PBS supplemented with 0.1% Tween. Blots were developed using Pierce ECL Western Blotting Substrate (Thermo Fisher Scientific), for 1 minutes at RT. Blots were imaged on a LAS4000 Imagequant (GE Healthcare).

### C5a formation assay

High density C3b beads, prepared as described above (C3b beads binding assay), were incubated with UNbC3b-1 (4.1 – 1000 nM), FB_DGF_ (20 µg/mL), FD (5 µg/ml), C3 (20 µg/mL) and/or C5 (20 µg/mL). Components were added subsequently, with in between a 10-minute incubation step at RT, while shaking. The exact sequence of adding components is indicated in each figure and corresponding legends. At the final incubation step, supernatants were collected and C3a and C5a release was measured using a calcium flux assay (9, 10). For this, U937 C3aR and C5aR cells(9), at a concentration of 5×10^6^ cells/ml, were incubated with 0.5 µM Fluo-3-AM ester (Invitrogen) in RPMI (Lonza) supplemented with 0.05% human serum albumin (HSA, Sanquin). Next, cells were diluted to a concentration of 1×10^6^ cells/ml and divided in 200 µL/sample. Next, 20 µl supernatant, containing the C3a or C5a stimulus, was added and fluorescence was measured in time using a FACS Verse flow cytometer (BD). To calculate the amount of C3a and C5a present in the supernatants, C3a and C5a standard curves were made using purified C3a and C5a.

### SEC: nanobody interaction with C3 and C3b

UNbC3b-1-a488 (2.3 µM) was incubated with C3 (2.3 µM), C3b (2.3 µM) or buffer (25 mM Hepes + 150 mM NaCl, pH 7.4) for 15 minutes, at RT. As controls, C3 andC3b, samples were incubated with buffer instead of UNbC3b-1. Next, samples were filtered with a 0.22 µm Costar® SpinX tube (Corning) and 50 µL sample was loaded on a Superose 6 Increase 3.2/300 column (GE Healthcare), using the AKTA-Explorer (GE Healthcare), which measured the OD_280_ and OD_488_.

### SPR affinity determination

Binding affinity of UNbC3b-1 for C3b was assessed using a White Fox fiber optic surface plasmon resonance (FO-SPR) sensor (Fox Biosystems). UNbC3b-1-biotin (330 nM) was immobilized on streptavidin FO-SPR probes (Fox Biosystems) in PBS, pH 7.4, supplemented with 0.01% Tween-20 (PBS-T_0.01%_) for 300 seconds, at 26 °C, while shaking at 1000 rpm. Next, probes were incubated with C3b (1.56 – 12.5 nM) in PBS-T_0,01%_, at 26°C, while shaking at 1000 rpm, to allow association. After 1800 seconds, probes were incubated with PBS-T_0,01%_ for 7200 seconds, at 26°C, while shaking at 1000 rpm, to allow dissociation. FO-SPR data were collected by FOx software (FOx Biosystems) and on- and off-rates were fitted using TraceDrawer, using a 1:1 model with global B_max_, global k_on_, global k_off_ and a constant BI.

### Cryogenic electron microscopy (Cryo-EM) sample preparation and data collection

Purified C3b was mixed with UNbC3b-1 at a 1:3 molar ratio and incubated on ice for 15 minutes. Following the incubation, 3.5 µl was applied to a glow discharged (Pelco) R 1.2/1.3 200 mesh Au holey carbon grid (Quantifoil) and plunge frozen using a Vitrobot Mark IV (Thermo Fisher Scientific, TFS) at 10°C and 80% relative humidity. Data was collected using a TFS Talos Arctica 200 keV cryogenic electron microscope equipped with a Gatan K2 direct electron detector and post column energy filter. 1568 movies were collected using SerialEM at a pixel size of 1.359 Å, energy filter slit width of 20 eV, and a total dose of 50 e-/Å.

### Cryo-EM data processing and model building

The resulting data was processed using the CryoSPARC v3.3/4.5.3 software suite (Fig **S6A**, (22)). Following import, the movies were motion corrected using Patch Motion Correction and CTF estimation was done using Patch CTF. Next, the Blob Picker was used to select an initial set of particles that were then extracted at a pixel size of 5.436 Å (Fourier cropped from 224 to 56 pix) and used to generate 2D classes. After two rounds of cleaning, five ab-initio models were generated from 253,740 particles. Class 3 was selected for further processing. The 76,465 particles were reextracted unbinned with a box size of 224 pix and these particles were subjected to homogeneous refinement using Class 3 ab-initio model as the initial starting volume. The resulting density map was further refined using non-uniform refinement (NU-refine, (42)), generating a final density map with global resolution of 4.19 Å. Subsequently, the crystal structure of C3b (PDB 2I07, (43)) rigid-body fit into the density map filtered to 5 Å using the Fit in Map function in ChimeraX 1.8 (44). This revealed an additional density in the region of the MG ring that could accommodate the UNbC3b-1 nanobody. Alphafold2 2.3.1 was used to generate an initial model of UNbC3b-1 using a Colabfold v1.5.1 notebook offered via Google Collaboratory (45, 46). Comparisons of the final C3b:UNbC3b-1 with other published proteins were made in ChimeraX using the Matchmaker function.

### Data analysis

Graphs were created with GraphPad Prism 9.3.0. Curves were fitted with GraphPad Prism 9.3.0. or with Tracedrawer 1.9.2. Nanobody sequences were analyzed and aligned using PipeBio.

## SUPPORTING INFORMATION

This article contains supporting information.

## FUNDING AND ADDITIONAL INFORMATION

This work was mainly funded by the Netherlands Organisation for Scientific Research (NWO) under the TTW Industrial Doctorate (grant agreement no. NWA.ID.17.036 to EMS). The project was also supported by the European Research Council (ERC) under the European Union’s Horizon 2020 Excellent Science programme (grant agreement No. 101001937, ERC-ACCENT to SHMR and grant agreement No. 787241, ERC-AdG to PG).

## ABBREVIATIONS

2YT: 2 times concentrated yeast extract tryptone medium
AF2: AlphaFold2
AMD: age-related macular degeneration
ANA: anaphylatoxin domain
AP: alternative pathway
bac-C3b_n_: bacteria opsonized with high densities of C3b molecules
BS3: bis(sulfosuccinimidyl)suberate
BSA: bovine serum albumin
C3(H_2_O): hydrolyzed form of C3
C345c/CTC: C-terminal domain
C3b-bio: C3b-PEG11-biotin
C3G: C3 glomerulopathy
CDR: complementarity determining region
CFU/ml: colony forming units per mL
CP: classical pathway
CUB: C1r/C1s, Uegf, Bmp1 domain
DBCO: dibenzocyclooctyne
DIG: digoxigenin
EM: wells that were not inoculated with bacteria
ERC: European research council
FB: factor B
FB_DGF_: FB double gain of function mutant
FB_WT_: FB wildtype
FD: factor D
FH: factor H
FI: factor I
FO-SPR: fiber optic surface plasmon resonance
HSA: human serum albumin
IPTG: isopropylthio-β-galactoside
IRR: nanobody with irrelevant target
KDO-N_3_: azide-modified keto-deoxy-octulosonate
LB: Luria-Bertani
LD: low density
LNK: linker domain
LP: lectin pathway
LPS: lipopolysaccharides
MAC: membrane attack complex
MG: macroglobulin
MH-clones: nanobodies containing a Myc-6×His-tag
NWO: Netherlands Organisation for Scientific Research
O/N: overnight
PBMC: peripheral blood mononuclear cells
PBS: phosphate buffered saline
PBS-T: PBS supplemented with 0.05% Tween-20
PBS-T_0.01%_: PBS supplemented with 0.01% Tween-20
PER: bacterium containing an empty vector
peri-Nbs: nanobodies in the crude periplasmic extracts
PFA: paraformaldehyde
PNH: paroxysmal nocturnal hemoglobinuria
QE19: polyclonal rabbit-anti-nanobody antibodies
QE19-a647: QE19 antibodies coupled with fluorophore Alexa 647
raE: rabbit erythrocytes
RT: room temperature
SEC: size exclusion chromatography
shE: sheep erythrocytes
shEA: sheep erythrocytes opsonized with antibodies (amboceptor)
SP: serine protease
SPR: surface plasmon resonance
TED: thioester-containing domain
UNbC3b-1-a488: nanobody UNbC3b-1 labeled with fluorophore Alexa 488
UNbC3b-1-bio: nanobody UNbC3b-1 biotinylated
UNbC3b-1: nanobody UNbC3b-1 with an LPETG-6×His tag
VBS: veronal buffered saline supplemented with 145 nM NaCl, at pH 7.4
VBS/Mg/EGTA: VBS supplemented with 5 mM MgCl_2_ and 10 mM EGTA
VBS/MgCl_2_/Tween: VBS supplemented with 2.5 mM MgCl_2_ and 0.05% Tween-20

