## Supplemental figures and legends for "Identification of a C3b-specific nanobody that does not bind C3 and blocks alternative pathway convertases"

SFig. 1

A

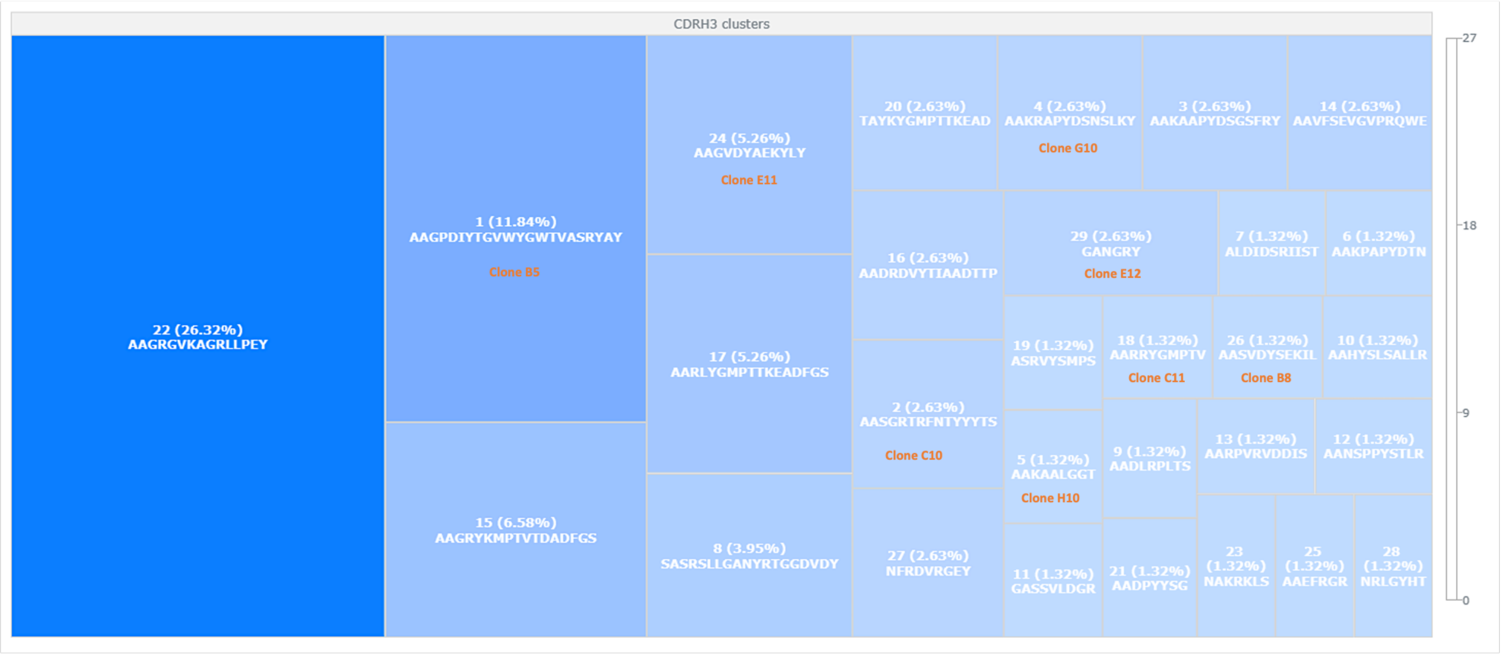

CDRH3 clusters
22 (26.32%)
AAGRGVKAGRLLPEY
1 (11.84%)
AAGPDIYTGVWYGWTVASRYAY
Clone B5
15 (6.58%)
AAGRYKMPTVTDADFGS
24 (5.26%)
AAGVDYAEKYLY
Clone E11
17 (5.26%)
AARLYGMPTTKEADFGS
8 (3.95%)
SASRSLLGANYRTGGDVDY
20 (2.63%)
TAYKYGMPTTKEAD
4 (2.63%)
AAKRAPYDSNSLKY
Clone G10
3 (2.63%)
AAKAAPYDSGSFRY
14 (2.63%)
AAVFSEVGVPRQWE
16 (2.63%)
AADRDVYTIAADTTP
29 (2.63%)
GANGRY
Clone E12
7 (1.32%)
ALDIDSRIIST
6 (1.32%)
AAKPAPYDTN
2 (2.63%)
AASGRTRFNTYYYTS
Clone C10
19 (1.32%)
ASRVYSMPS
18 (1.32%)
AARRYGMPTV
Clone C11
26 (1.32%)
AASVDYSEKIL
Clone B8
10 (1.32%)
AAHYSLSALLR
5 (1.32%)
AAKAALGGT
Clone H10
9 (1.32%)
AADLRPLTS
13 (1.32%)
AARPVRVDDIS
12 (1.32%)
AANSPPYSTLR
27 (2.63%)
NFRDVRGEY
11 (1.32%)
GASSVLDGR
21 (1.32%)
AADPYYSG
23
(1.32%)
NAKRKLS
25
(1.32%)
AAEFRGR
28
(1.32%)
NRLGYHT
27
18
9
0


B

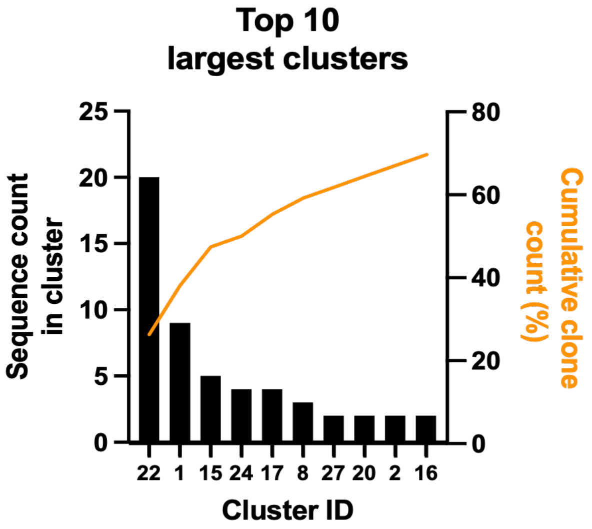

Top 10
largest clusters
Sequence count
in cluster
25
20
15
10
5
0
80
60
40
20
0
Cumulative clone
count (%)
22 1 15 24 17 8 27 20 2 16
Cluster ID


C

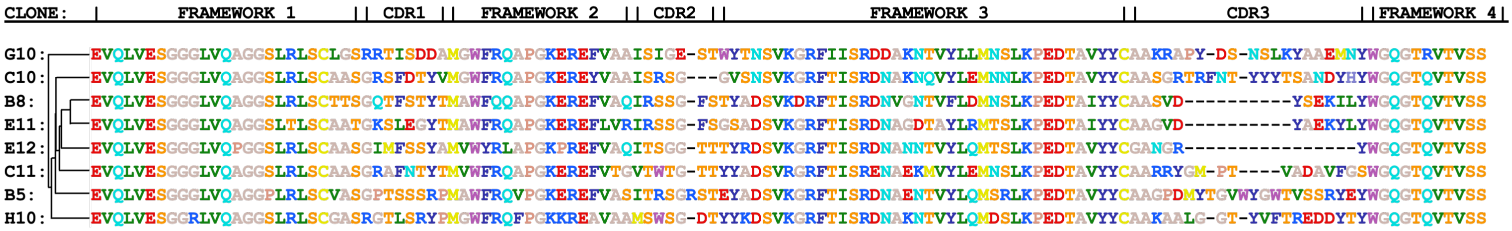

CLONE: | FRAMEWORK 1 || CDR1 || FRAMEWORK 2 || CDR2 || FRAMEWORK 3 || CDR3 || FRAMEWORK 4 |
G10: EVQLVESGGGLVQAGGSLRLSCLGSRRTISDDAMGWFRQAPGKEREFVAAISIGE-STWYTNSVKGRFIISRDDAKNTVYLLMNSLKPEDTAVYYCAAKRAPY-DS-NSLKYAAEMNYWGQGTRVTVSS
C10: EVQLVESGGGLVQAGGSLRLSCAASGRSFDTYVMGWFRQAPGKEREYVAAISRSG---GVSNSVKGRFTISRDNAKNQVYLEMNNLKPEDTAVYYCAASGRTRFNT-YYYTSANDYHYWGQGTQVTVSS
B8: EVQLVESGGGLVQAGGSLRLSCTTSGQTFSTYTMAWFQQAPGKEREFVAQIRSSG-FSTYADSVKDRFTISRDNVGNTVFLDMNSLKPEDTAIYYCAASVD----------YSEKILYWGQGTQVTVSS
E11: EVQLVESGGGLVQAGGSLTLSCAATGKSLEGYTMAWFRQAPGKEREFLVRIRSSG-FSGSADSVKGRFTISRDNAGDTAYLRMTSLKPEDTAIYYCAAGVD----------YAEKYLYWGQGTQVTVSS
E12: EVQLVESGGGLVQPGGSLRLSCAASGIMFSSYAMVWYRLAPGKPREFVAQITSGG-TTTYRDSVKGRFTISRDNANNTVYLQMTSLKPEDTAVYYCGANGR----------------YWGQGTQVTVSS
C11: EVQLVESGGGLVQAGGSLRLSCAASGRAFNTYTMVWFRQAPGKEREFVTGVTWTG-TTYYADSVRGRFTISRENAEKMVYLEMNSLKPEDTAVYYCAARRYGM-PT----VADAVFGSWGQGTQVTVSS
B5: EVQLVESGGGLVQAGGPLRLSCVASGPTSSSRPMAWFRQVPGKEREFVASITRSGRSTEYADSVKGRFTISRDNAENTVYLQMSRLKPEDTAVYYCAAGPDMYTGVWYGWTVSSRYEYWGQGTQVTVSS
H10: EVQLVESGGRLVQAGGSLRLSCGASRGTLSRYPMGWFRQFPGKKREAVAAMSWSG-DTYYKDSVKGRFTISRDNAKNTVYLQMDSLKPEDTAVYYCAAKAALG-GT-YVFTREDDYTYWGQGTQVTVSS

SFig. 2

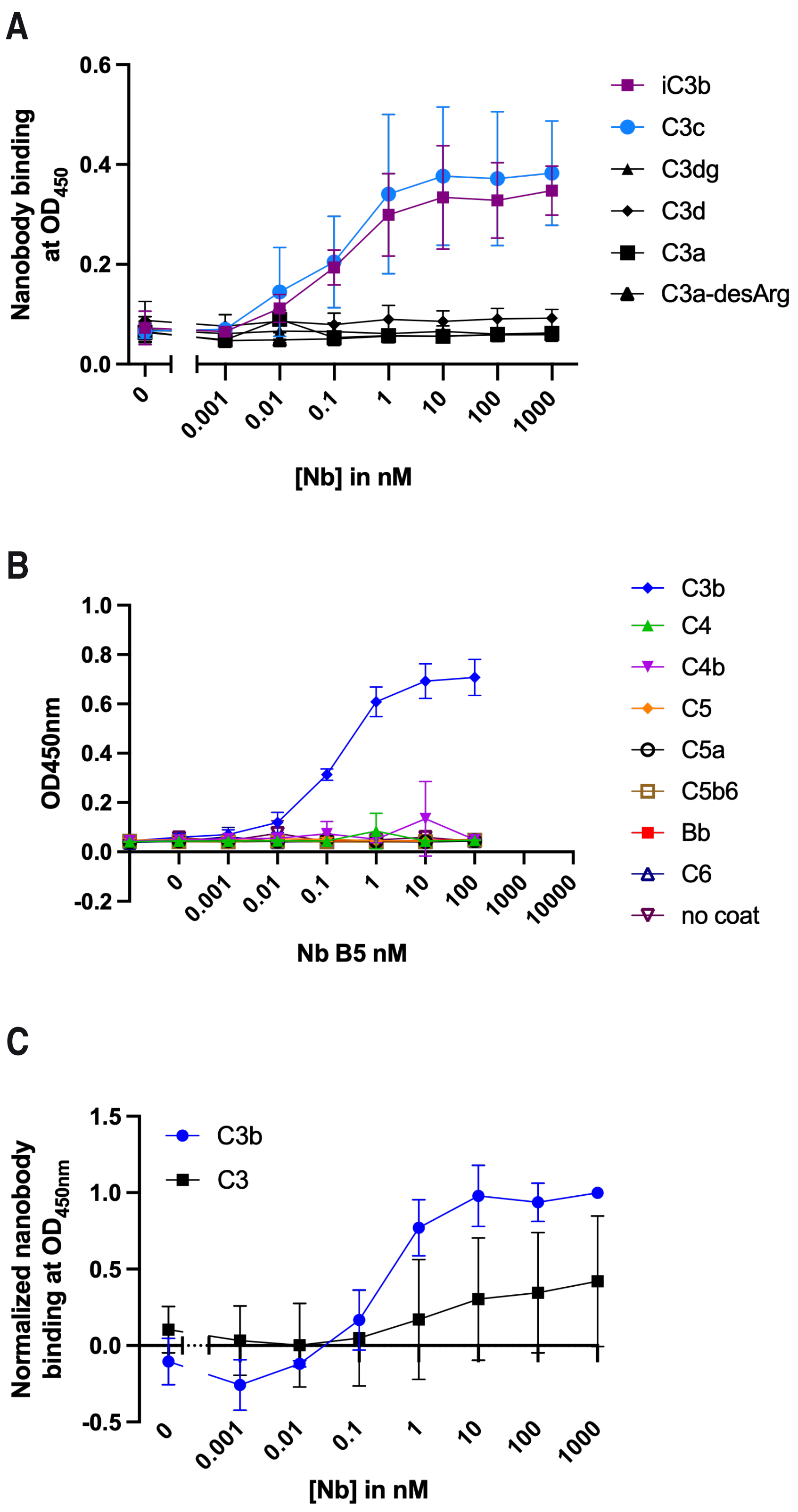
A
Nanobody binding at $OD_{450}$
0.6
0.4
0.2
0.0
0
0.001
0.01
0.1
1
10
100
1000
[Nb] in nM
iC3b
C3c
C3dg
C3d
C3a
C3a-desArg
B
OD450nm
1.0
0.8
0.6
0.4
0.2
0.0
-0.2
0
0.001
0.01
0.1
1
10
100
1000
10000
Nb B5 nM
C3b
C4
C4b
C5
C5a
C5b6
Bb
C6
no coat
C
Normalized nanobody binding at $OD_{450nm}$
1.5
1.0
0.5
0.0
-0.5
0
0.001
0.01
0.1
1
10
100
1000
[Nb] in nM
C3b
C3

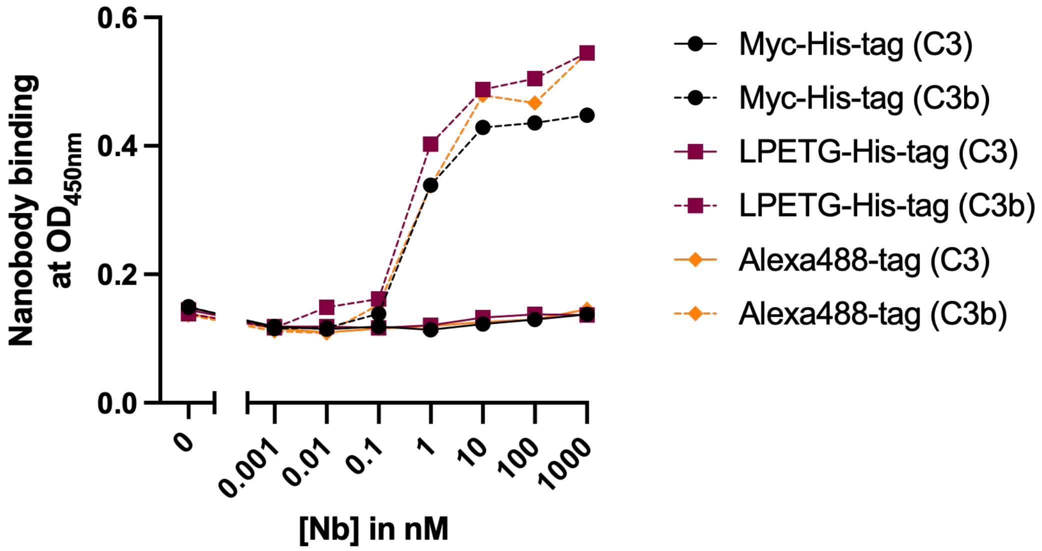
Nanobody binding at $OD_{450nm}$
0.6
0.4
0.2
0.0
0
0.001
0.01
0.1
1
10
100
1000
[Nb] in nM
Myc-His-tag (C3)
Myc-His-tag (C3b)
LPETG-His-tag (C3)
LPETG-His-tag (C3b)
Alexa488-tag (C3)
Alexa488-tag (C3b)

A

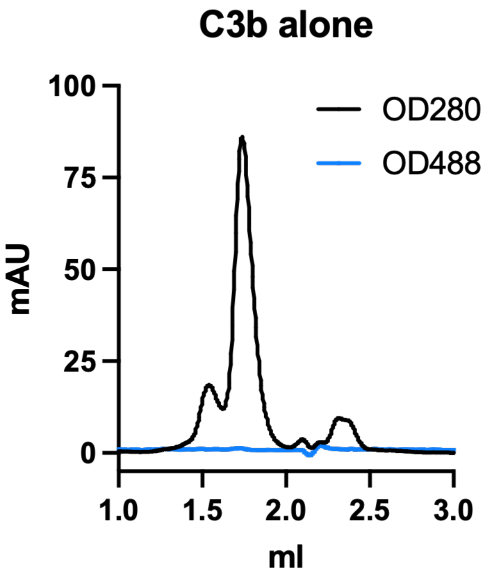
C3b alone
100
75
50
25
0
mAU
1.0
1.5
2.0
2.5
3.0
ml
OD280
OD488

B

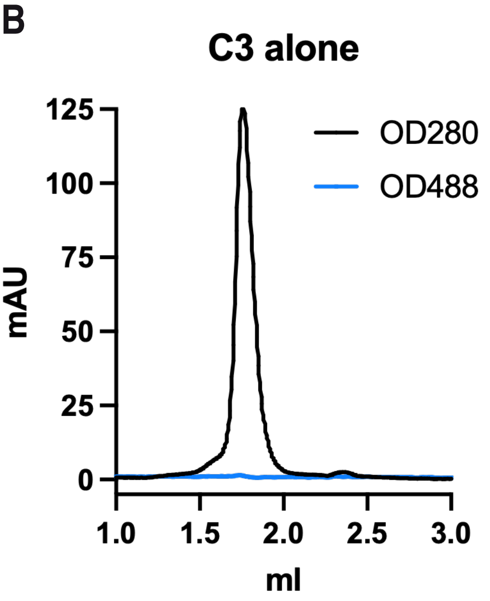
C3 alone
125
100
75
50
25
0
mAU
1.0
1.5
2.0
2.5
3.0
ml
OD280
OD488

C

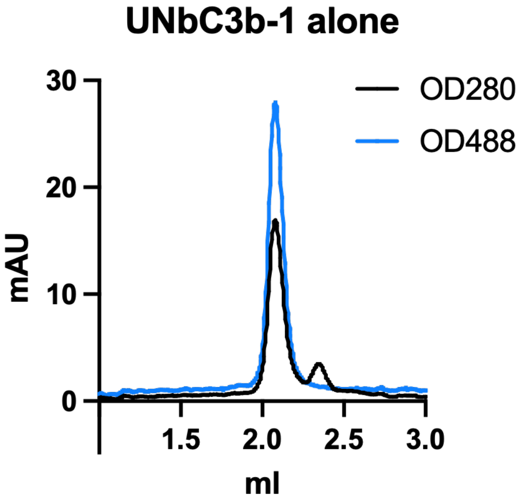
UNbC3b-1 alone
30
20
10
0
mAU
1.5
2.0
2.5
3.0
ml
OD280
OD488

SFig. 5

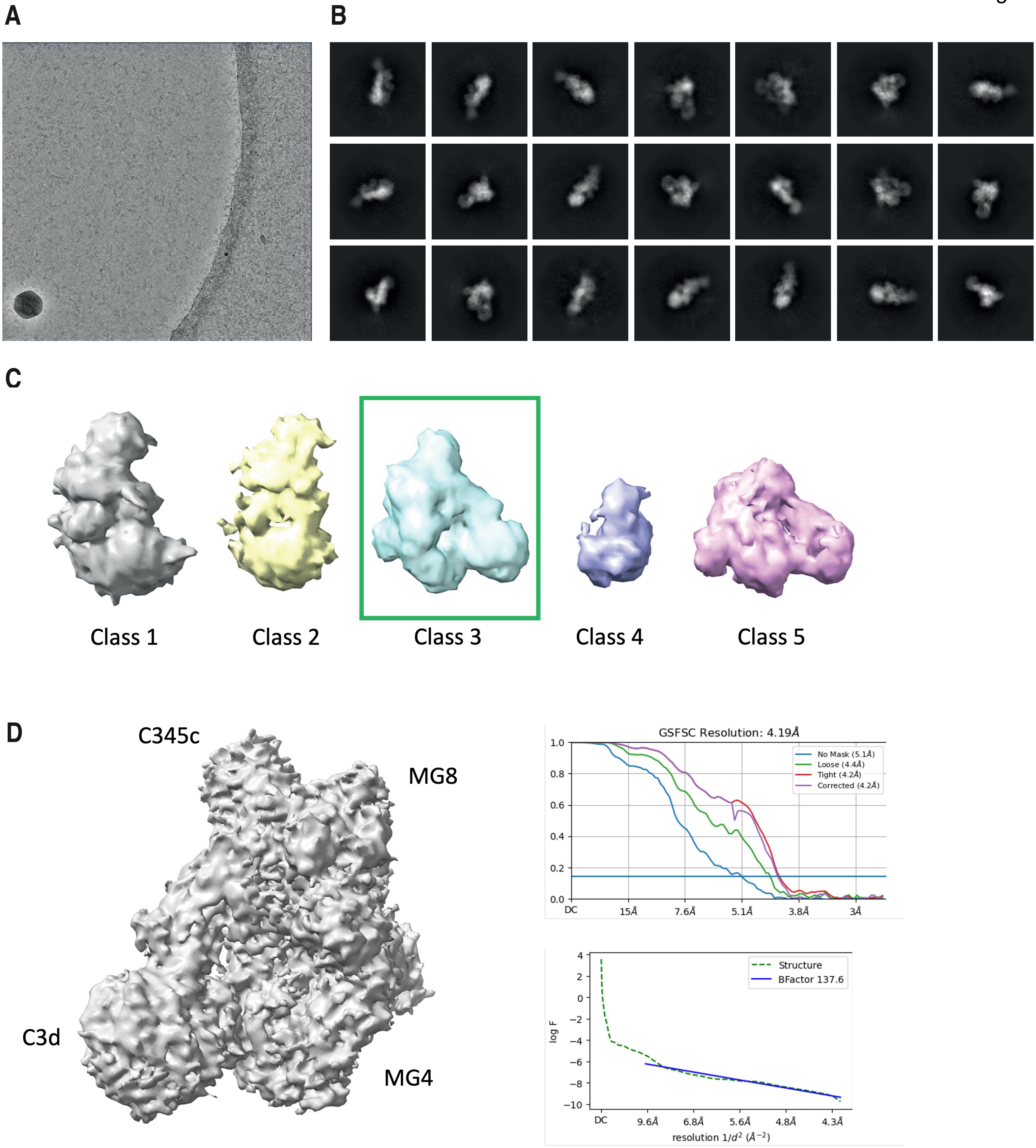

A
B
C
Class 1
Class 2
Class 3
Class 4
Class 5
D
C345c
MG8
C3d
MG4
GSFSC Resolution: 4.19Å
No Mask (5.1Å)
Loose (4.4Å)
Tight (4.2Å)
Corrected (4.2Å)
DC
15Å
7.6Å
5.1Å
3.8Å
3Å
Structure
BFactor 137.6
log F
9.6Å
6.8Å
5.6Å
4.8Å
4.3Å
resolution 1/d² (Å⁻²)

A

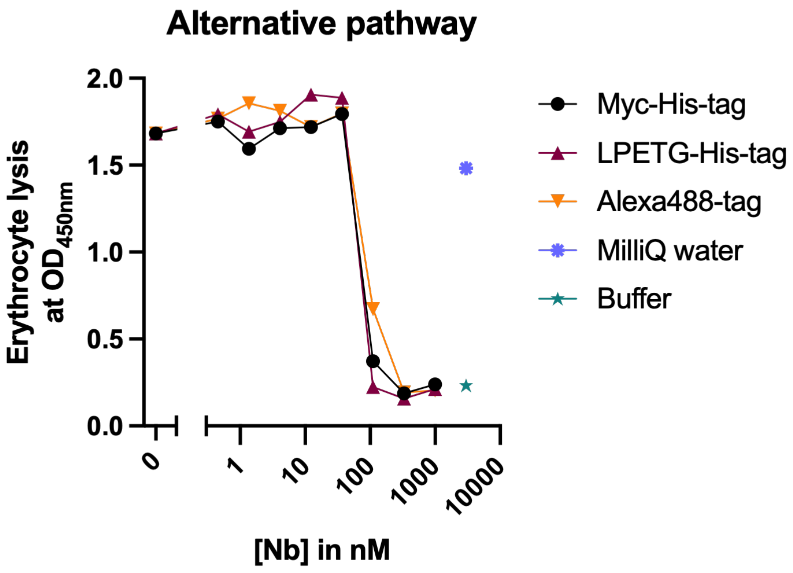
Alternative pathway
Erythrocyte lysis at OD450nm
2.0
1.5
1.0
0.5
0.0
0
1
10
100
1000
10000
[Nb] in nM
Myc-His-tag
LPETG-His-tag
Alexa488-tag
MilliQ water
Buffer

B

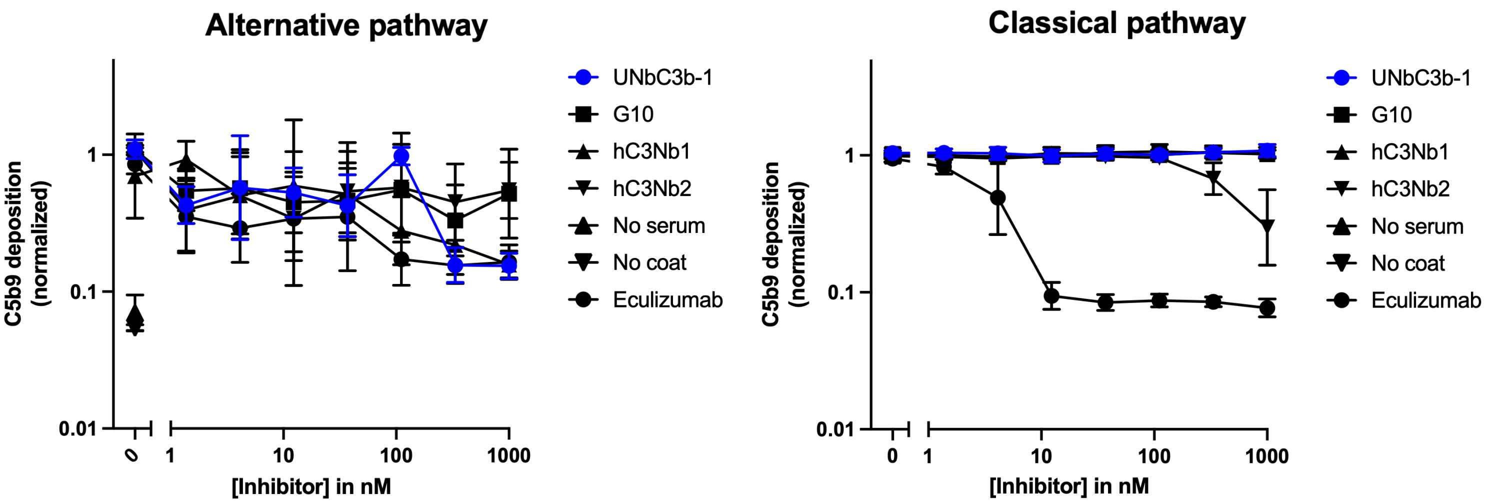
Alternative pathway
C5b9 deposition (normalized)
1
0.1
0.01
0
1
10
100
1000
[Inhibitor] in nM
UNbC3b-1
G10
hC3Nb1
hC3Nb2
No serum
No coat
Eculizumab
Classical pathway
C5b9 deposition (normalized)
1
0.1
0.01
0
1
10
100
1000
[Inhibitor] in nM
UNbC3b-1
G10
hC3Nb1
hC3Nb2
No serum
No coat
Eculizumab

**Supporting Figure 1: Sequence analysis of all screened clones.** (A) Schematic representation of different clusters containing >80% sequence similarity in the CDR-H3 region. CDR-H3 amino acid sequences are depicted, and cluster sizes are indicated in percentages and color (the darker the higher the percentage). The cluster origin of the eight selected nanobodies is annotated in orange text. (B) The cluster size of the top 10 largest clusters is depicted in a bar-graph with the cumulative cluster count depicted in orange. (C) Amino acid sequence alignment of the 8 selected clones, with the different framework and CDR regions indicated. Phylogenetic tree indicates cluster relatedness and colors depict types of amino acid, using the shapely color scheme.

**Supporting figure 2: UNbC3b-1 specifically binds to C3b, and not to other complement components.** (A) ELISA with UNbC3b-1 binding to different C3 fragments coated on microtiter plates. Curves of iC3b (purple) and C3c (blue) are colored to highlight that UNbC3b-1 binds to these fragments. (B) Cross reactivity of UNbC3b-1 with other complement components. Bb, C4, C4b, C5, C5a, C5b6, and C6 were coated on microtiter plates and different concentrations of UNbC3b-1-biotin were added. Nanobody binding was detected with streptavidin-PO. (C) ELISA to measure UNbC3b-1 binding to C3b and C3 coated microtiter plates. (A, C) nanobody binding was assessed using polyclonal rabbit-anti-VHH QE19 antibodies and donkey-anti-rabbit-HRP antibodies, at an OD of 450 nm. Data represent mean ± SD of two (A) or three (B, C) individual experiments.

**Supporting Figure 3: Myc/LPETG-His and fluorophore tags do not influence UNbC3b-1 binding to C3b.** ELISA with C3 and C3b coated on microtiter plates which were incubated with different batches of nanobody, in different concentrations. Nanobody binding was assessed using polyclonal rabbit-anti-VHH QE19 antibodies and donkey-anti-rabbit-HRP antibodies, at an OD of 450 nm.

**Supporting figure 4: SEC runs of C3b, C3, and UNbC3b-1.** (A-C) Size exclusion chromatography controls with C3b (A), C3 (B), and UNbC3b-1-a488 (C) run individually on a Superose 6 Increase column. The OD was measured at 280 nm and 480 nm. Data represent two individual experiments.

**Supporting figure 5: Cryo-EM image processing pipeline for determination of UNbC3b-1 binding site.** (A) Sample micrograph showing protein distribution. (B) Representative set of 2D classes used for particle cleaning. (C) Five ab-initio models were generated from 253,740 particles, identifying one class that had the characteristics of C3b (green box). (D) Further processing of class 3 (76,465 particles) resulted in the in final electron density map (left) with a global resolution of 4.19 Å, and the corresponding final resolution FSC curve and B-factor plots (right). This map was subsequently used for modeling.

**Supporting figure 6: Inhibition of complement by (tagged) nanobodies.** (A) AP-mediated hemolysis of rabbit erythrocytes incubated with 10% human serum and different batches of nanobody in different concentrations. MilliQ and buffer were taken along as controls for 100% and 0% lysis, respectively. Data presented are of one individual experiment. (B) Left: AP activation on an LPS-coated microtiter plate, incubated with 10% human serum and different concentrations of inhibitors. Right: CP activation on an IgM-coated microtiter plate, incubated with 2.5% human serum and different concentrations of inhibitors. Deposition of complement activation product C5b-9 on the plate was measured using the mouse-anti-aE11 and goat-anti-mouse-HRP antibodies, at OD450. Data points were normalized to the condition without inhibitor and graphs represent mean ± SD of three individual experiments.
